# Dissecting the initiation of female meiosis in the mouse at single-cell resolution

**DOI:** 10.1101/803668

**Authors:** Wei Ge, Jun-Jie Wang, Rui-Qian Zhang, Shao-Jing Tan, Fa-Li Zhang, Wen-Xiang Liu, Lan Li, Xiao-Feng Sun, Shun-Feng Cheng, Paul W. Dyce, Massimo De Felici, Wei Shen

## Abstract

Germ cell meiosis is one of the most finely orchestrated events during gametogenesis with distinct developmental patterns in males and females. However, in mammals, the molecular mechanisms involved in this process remain not well known. Here, we report detailed transcriptome analyses of cell populations present in the mouse female gonadal ridges (E11.5) and the embryonic ovaries from E12.5 to E14.5 using single cell RNA sequencing (scRNA seq). These periods correspond with the initiation and progression of meiosis throughout the first stage of prophase I. We identified 13 transcriptionally distinct cell populations and 7 transcriptionally distinct germ cell subclusters that correspond to mitotic (3 clusters) and meiotic (4 clusters) germ cells. By comparing the signature gene expression pattern of 4 meiotic germ cell clusters, we found that the 4 cell clusters correspond to different cell status en route to meiosis progression, and therefore, our research here characterized detailed transcriptome dynamics during meiotic prophase I. Reconstructing the progression of meiosis along pseudotime, we identified several new genes and molecular pathways with potential critical roles in the mitosis/meiosis transition and early meiotic progression. Last, the heterogeneity within somatic cell populations was also discussed and different cellular states were identified. Our scRNA seq analysis here represents a new important resource for deciphering the molecular pathways driving meiosis initiation and progression in female germ cells and ovarian somatic cells.

## INTRODUCTION

In mammals, the primordial germ cells (PGCs) are considered germline stem cells which give rise to the male and female gametes, ultimately responsible for the survival of species and the transmission of genetic information across generations (Cinalli et al., 2008, Trapnell et al., 2014). A central process of which is meiosis. Meiosis in female mammals begins in the developing ovaries during the embryonic/fetal period arresting at the end of prophase I around or just after birth (De Felici, 2016). In mice, female PGCs colonize the gonadal ridges from E10.5 to E12.5 and after some rounds of mitotic division begin to enter meiosis between E13.5 and E14.5 (Grive & Freiman, 2015, Wang et al., 2017). During the mitosis-meiosis transition, germ cells undergo extensive changes in gene expression as they progress through the leptotene, zygotene, pachytene and diplotene stages until arresting at the dictyate stage of meiosis prophase I (Wang et al., 2017). Meiosis initiation has long been considered as the gatekeeper of successful gametogenesis (Ge et al., 2015a). In fact, the fidelity of meiosis initiation along with progression is vital for future reproductive health and defects in such processes can lead to reproductive diseases including premature ovarian failure, polycystic ovary syndrome and even infertility (Mandon-Pepin et al., 2008, Qin et al., 2015, Tucker et al., 2016). However, in mammals due to the paucity of information regarding the molecular mechanisms regulating the initiation and progression of meiosis, the aetiology of such reproductive diseases remains elusive. In order to obtain insights into these processes, researchers have used a variety of experimental approaches. For example, in the mouse, many attempts have been performed using the *in vitro* culture of embryonic gonads and isolated PGCs and more recently in producing gametes from various stem cell types to reproduce meiotic entry and progression (Ge et al., 2015a, Saitou & Miyauchi, 2016). For obvious reasons, this latter approach is particularly useful in humans where promising results have been very recently obtained (Yamashiro et al., 2018). Several studies have reported the successful generation of germ cell-like cells from stem cells, however, until recently convincing evidence that such cells were able to correctly enter meiosis were lacking (Handel et al., 2014, Sun et al., 2014, Tedesco et al., 2011). In 2016, Zhou’s group and Hayashi’s group, reported the successful derivation of functional sperm and oocytes from pluripotent stem cells, respectively (Hikabe et al., 2016, Zhou et al., 2016). Although these studies provide valuable sources for investigating mammalian meiosis under complete *in vitro* conditions, the germ cell differentiation efficiency was limited, and many germ cells showed abnormal meiotic entry as evidenced by the increased percentage of asynapsis in comparison to endogenous and *in vitro* cultured germ cells.

After many decades of work identifying compounds able to induce PGC formation in females or inhibit male PGC entry into meiosis, a retinoic acid (RA)-*Stra8* signaling pathway has emerged as a key regulator of meiosis initiation in mice (Bowles & Koopman, 2007, Griswold et al., 2012). Today, it is widely accepted that in mammals, RA secreted from the mesonephroi activates meiotic gene expression (i.e., *Sycp3*, *Sycp1*, *Stra8* and *Rec8*) and initiates meiotic programs in female PGCs (Griswold et al., 2012). However, recent studies from Miyauchi et al., in accord with previous results by Farini et al., found that RA alone is not sufficient to initiate meiosis onset in PGC-like cells produced from stem cells *in vitro* and demonstrated that the crosstalk between RA signaling and BMP signaling is pivotal for the activation of meiotic transcriptional cascades (Farini et al., 2005, Miyauchi et al., 2017). Previous work has largely focused on the initiation of meiosis while key regulators of meiotic progression remain to be identified.

In the present study, by using high-throughput single-cell RNA sequencing (scRNA seq), we analyzed the transcriptome data from 19,387 individual cells that were obtained from gonadal ridges and ovaries of E11.5 - E14.5 mouse embryos. Based on these data, we successfully identified germ cell and somatic cell subpopulations, furthermore, we characterized detailed germ cell transcriptome gene expression signatures en route to meiosis. Pseudotime ordering analysis successfully recapitulated germ cell meiosis initiation trajectory and revealed key molecular events involved during the transition from mitosis-meiosis. We also discussed the heterogeneity of the ovarian somatic cells. The results obtained here provide novel information and insights into the initiation and progression of meiosis and the diversity of somatic cells and lineages within the developing gonads.

## RESULTS

### Identification and characterization of the ovarian cell populations

To decipher the gene expression landscape and dissect the cellular heterogeneity during the initiation of meiosis in female germ cells, in mice, we dissociated ovarian tissues from E11.5, E12.5, E13.5, and E14.5 embryos and prepared single cell suspensions for scRNA seq (Figure 1a and Supplementary Figure 1a) (Baltus et al., 2006, Koubova et al., 2006). To verify the *bona fide* progression of meiosis, we performed ovarian tissue cytospreads for SYCP3 and γH2AX to characterize the meiotic stage (Figure 1b) (Feng et al., 2014) and calculated the percentage of meiotic cells (SCP3 positive oocytes) at each time point (Figure 1c). Consistent with previous findings, germ cells in E11.5 and E12.5 were mitotic, and it was not until E13.5 that some germ cells entered meiosis (Kocer et al., 2009).

**Figure 1.**
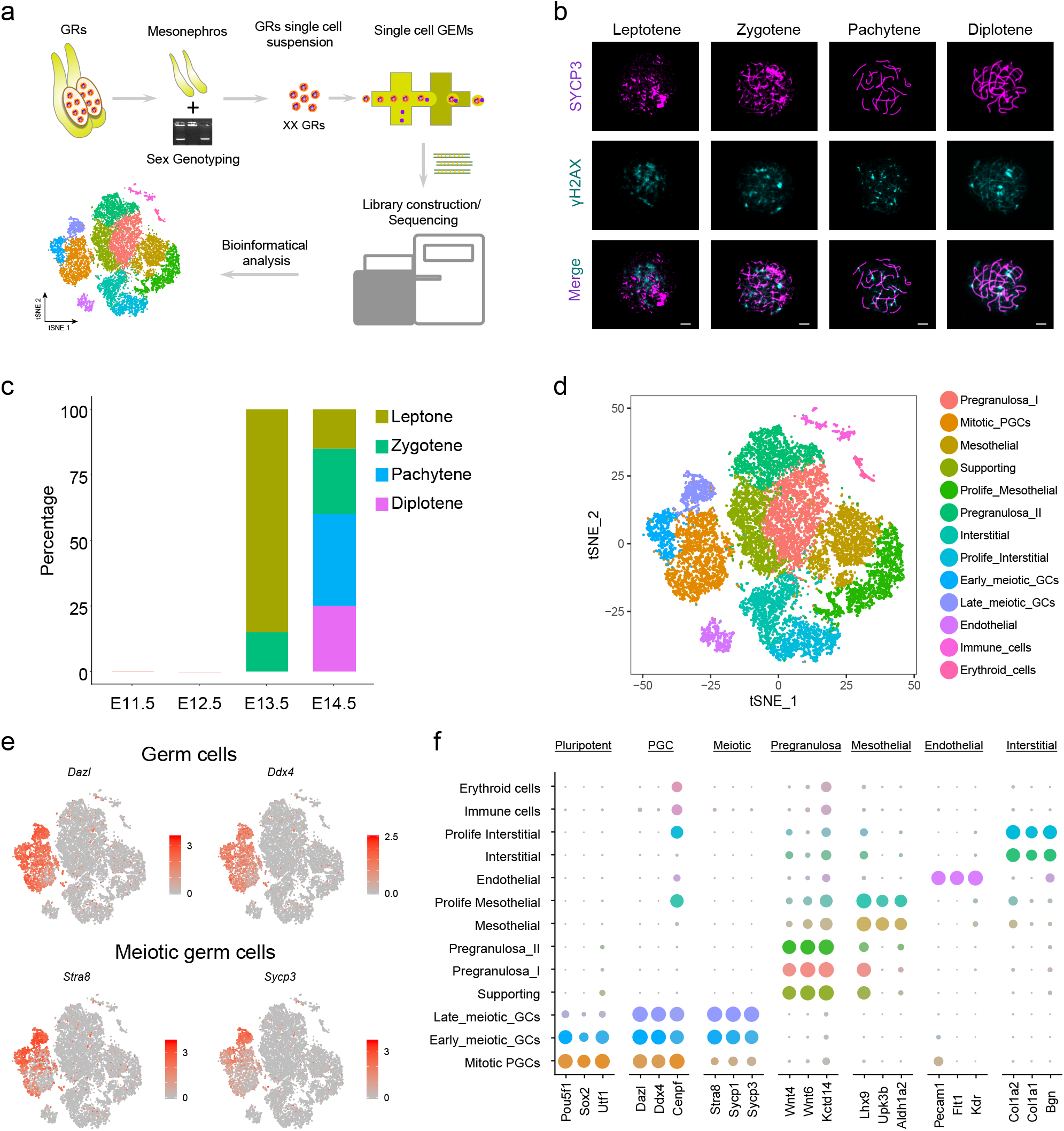
Experimental design and characterization of single cell clusters. **a** Diagram of sample preparation for scRNA seq. **b** Synaptonemal complex analysis of meiotic progression and representative SCP3 staining pattern at different stages (leptotene, zygotene, pachytene, and diplotene). Scale bars = 7.5 μm. **c** The percentage of premeiotic, leptotene, zygotene, pachytene, and diplotene stage germ cells at different developmental time-point (E11.5, E12.5, E13.5, E14.5). SCP3 positive oocytes emerged at E13.5 with ∼ 85 % oocytes at the leptotene stage and ∼ 15 % oocytes at the zygotene stage in C57BL/6 mice. When the fetus reached E14.5 oocytes at the pachytene and diplotene stages were first detected (accounting for ∼ 60 % of total meiotic oocytes). **d** tSNE plot labeled with cell identities. tSNE clustering identified 13 cell clusters in the developing gonads, and each cell type was labeled with a different color. **e** Marker gene expression projected onto the tSNE plot. The marker genes used were: *Dazl*, *Ddx4* for germ cells; *Stra8*, *Sycp3* for meiotic germ cells. **f** Dot plot of the pluripotent, meiotic germ cell, pregranulosa cell, mesothelial cell, endothelial cell and interstitial cell marker expression across all cell types. Each cell cluster was color-coded with their identity. The dot size represents the percentage of cells expressed the indicated genes in each cluster and the dot color intensity represents the average expression level of the indicated genes.

After filtering low quality cells based on the number of genes, unique molecular identifiers and the percentage of mitochondria genes (Supplementary Figure 1b), we obtained a total of 19,387 ovarian cells (4,916 cells for E11.5, 4,842 cells for E12.5, 4,842 cells for E13.5, and 4,787 cells for E14.5, respectively) and 19,321 genes, with the median genes per cell ranging from 3.097 ∼ 3,613 (Supplementary Figure 1a). To characterize cell identity, we integrated the four samples and performed t-distributed Stochastic Neighbor Embedding (tSNE) clustering analysis to dissect cellular heterogeneity among the ovarian cell populations (Figure 1d). After the tSNE projection, the four datasets integrated according to Seurat recommended algorithm (Supplementary Figure 1c). Further analysis delineated 13 transcriptionally distinct cell clusters across the four time points (Figure 1d). To identify germ cell populations within the plot, we visualized two genes that serve as classical germ cell markers, *Dazl* and *Ddx4*. Their expression across all single cells was determined and it was found that three cell clusters highly expressed the germ cell marker genes (Figure 1e) (Haston et al., 2009, Woods & Tilly, 2013). Pregranulosa cells or their precursors (supporting somatic cells) were identified within three distinct clusters on the basis of the elevated expression of *Wnt4* and *Wnt6* (Supplementary Figure 1d) (Jameson et al., 2012, McLaren, 2000). We also identified 5 ovarian somatic cells with their classic makers including mesothelial cells (*Lhx9* and *Upk3b*) (Kanamori-Katayama et al., 2011, Mazaud et al., 2002), interstitial cells (*Col1a2* and *Bgn*) (Piprek et al., 2018), endothelial cells (*Pecam1* and *Kdr*) (Brennan et al., 2002, Jeays-Ward et al., 2003) and two contaminative somatic cell populations from blood, immune cells (*Cd52* and *Car2*) and erythroid cells (*Alas2* and *Alad*) (Harigae et al., 2003, Lummertz da Rocha et al., 2018) (Supplementary Figure 1d). Thereby providing evidence of the heterogeneity of all somatic cell populations during the time frame analysed.

After this initial cluster identification, we began to analyze the dynamics of each cell population, principally of the germ cells and pregranulosa cells. To deconstruct the heterogeneous composition of the three germ cell clusters in the tSNE plot, we visualized the expression of the early PGC marker genes (*Pou5f1*, *Sox2*, *Utf1* and *Sall4*) (de Jong et al., 2008, Ohta et al., 2017), and the meiotic related genes (early: *Stra8*, *Scp3* and *Rec8*; late: *Scp1*, *Tex14* and *Mael*) (Figure 1e and Supplementary Figure 1d) (Malki et al., 2014, Morohaku et al., 2016, Watanabe & Nurse, 1999). These sets of genes showed distinct expression patterns: early PGC markers were mainly expressed in the lower part of clusters while the meiotic related genes were found mainly in the upper region. On this basis, we allocated premeiotic PGCs, along with early and late meiotic germ cells within the three germ cell clusters (Figure 1d). For the pregranulosa cell population, due to the lack of markers distinguishing supporting and pregranulosa cells, we interpreted such identities by combining the three *Wnt4* and *Wnt6* positive clusters with those of Supplementary Figure 1c. In particular, since in the mouse the majority of supporting cells differentiate into pregranulosa cells after gonadal sex determination (McLaren, 2000), we identified the supporting cell cluster representing the highest percentage of cells in the E11.5-E12.5 gonads along with two pregranulosa cell populations mainly in E13.5-E14.5 ovaries (Figure 1d). These analyses together with the calculation of the percentage of the cells within each cluster (Supplementary Figure 2a), preliminarily delineated the beginning and progression of meiosis in germ cells and the dynamics of the ovarian cell lineages.

To gain in-depth insight into the cluster-specific gene signatures and the functional gene categories of the ovarian cell populations, we also analyzed cell cluster-specific gene expression in more detail (Figure 1f and Supplementary Figure 2b). As expected, genes related to pluripotency, like *Pou5f1*, *Sox2* and *Utf1*, showed high expression in the early premeiotic PGCs that gradually decreased with the beginning and progression of meiosis. Conversely, the expression of meiotic genes such as *Stra8*, *Sycp1* and *Sycp3* significantly increased with the progression of meiosis (Figure 1f and Supplementary Figure 2c). To further validate our Seurat identified signature genes, we also performed immunofluorescence analysis on Seurat identified mitotic marker expression UTF1, which also showed similar results (Supplementary Figure 2d).

In total, we found 634, 654 and 1,189 differentially expressed genes (DEGs) in mitotic, early and late meiotic germ cells, respectively (Supplementary Table 1). Interestingly, gene ontology (GO) analysis of mitotic germ cells enriched mainly in pluripotent and cell cycle-related genes such as *Dppa5a*, *Utf1*, *Cenpf* and *Ifitm3*, and enriched GO terms of “cell division, ribonucleoprotein complex biogenesis and regulation of DNA metabolic process”. This supports the notion of active PGC proliferation prior to the mitosis-meiosis transition (De Felici & Farini, 2012). Early meiotic germ cells were enriched in *Stra8*, *Dazl*, *Smc1b*, *Hells* and *Sycp1*, while late meiotic germ cells enriched in *Smc1b*, *Sycp3*, *Sycp1*, *Tex101* and *Tex15*. Noteworthy, both early meiotic germ cells and late meiotic germ cells enriched the GO terms of “meiotic cell cycle, male meiotic nuclear division, and cellular response to DNA damage stimulus”, further confirming the entering into of meiosis at these stages (Supplementary Figure 2e). Among the somatic cells, granulosa cell lineage was mainly enriched in GO terms of “aldehyde biosynthetic process, reproductive structure development, and cellular response to extracellular stimulus” (Supplementary Figure 2f). Meanwhile, the interstitial and mesothelial populations were both similarly enriched in the cell cycle-related GO terms, such as “regulation of mitotic cell cycle” and “mitotic cell cycle” (Supplementary Figure 2g, 2h). Besides, interstitial cells enriched genes involved in “blood vessel development” and mesothelial cells enriched genes involved in “respiratory system development”. Regarding endothelial cells, our analysis revealed that GO terms of “angiogenesis, endothelial cell migration, and endothelial cell proliferation” were enriched, further confirming their endothelial identity (Supplementary Figure 2i).

### High-resolution dissection of germ cell meiotic progression at single-cell resolution

To dissect germ cell meiotic progress at a higher resolution, we subclustered the three germ cell clusters and re-performed tSNE projection (Figure 2a and 2b). Noteworthy, cell clustering using linear combination of the principal component analysis (PCA) algorithm also distinguished three germ cell clusters identified by tSNE into distinct populations, further emphasizing the different cellular states during the progression of meiosis (Figure 2a, right panel). More importantly, tSNE projection of all germ cells revealed 7 transcriptionally distinct subclusters (Figure 2b). Similar to the results reported in Figure 1d, we could also allocate the germ cells from E11.5 and E12.5 gonads mainly at the top of the tSNE plot and those from E13.5 and E14.5 at the middle and bottom of the tSNE plot, deciphering their entering and progression into meiosis (Figure 2b, left panel). Interestingly, by analyzing the top 5 cluster-specific expressed genes, we found that cluster 0 and 1 showed similar gene expression pattern, characterized by high expression of early meiotic markers *Stra8*, *Smc1b* and *Rec8*, while cluster 2, 3, 4 and 5 showed similar gene expression pattern, with high expression of *Dusp9*, *Wdr89*, and *Dppa5a* (Supplementary Figure 3a). For cluster 6, they showed high levels of late meiotic markers *Sycp3*, *Tex101*, and *Taf7l*. Taken together, the preliminary analysis demonstrated that the different germ cell clusters may represent different cell stages en route to meiosis.

**Figure 2.**
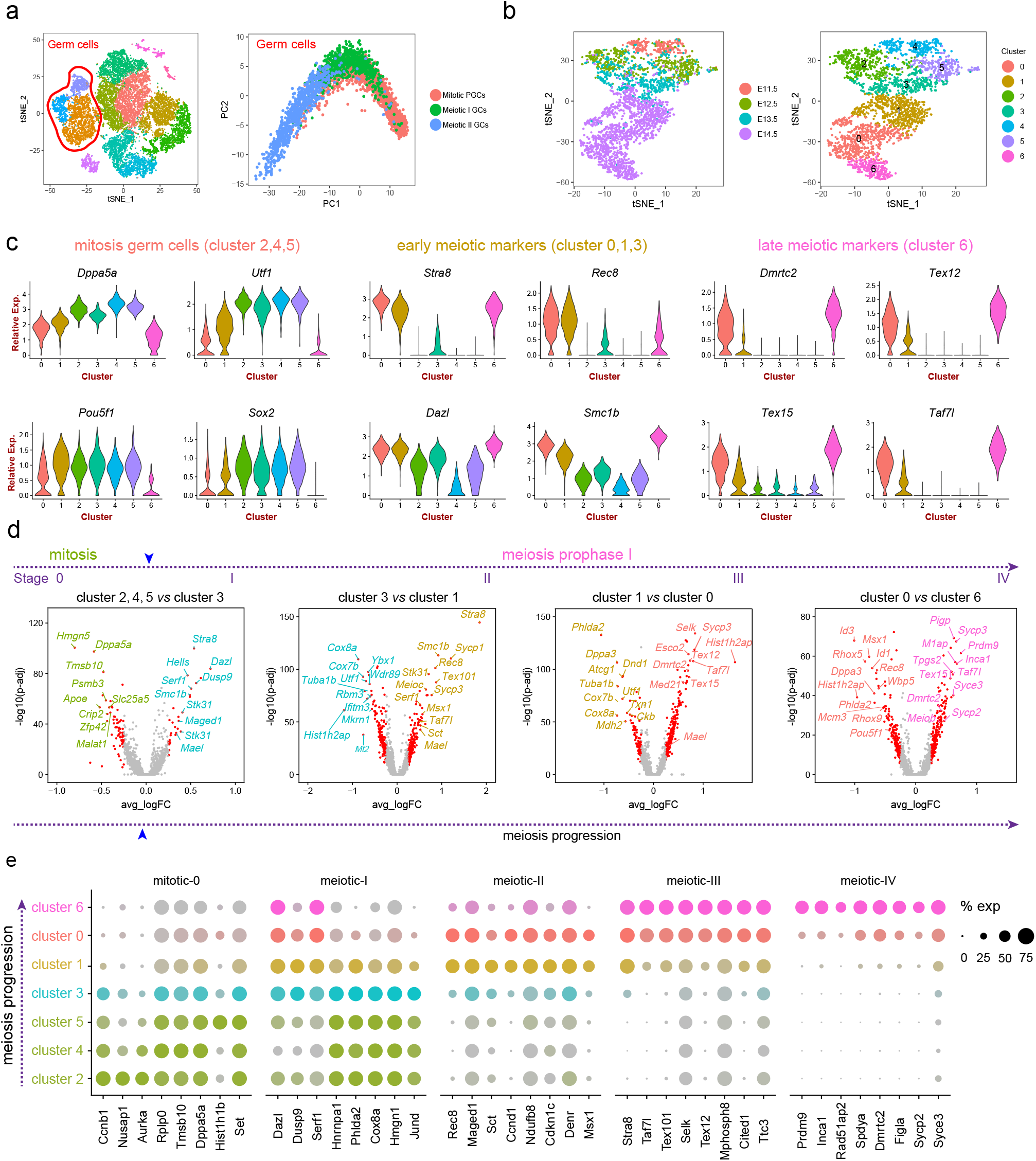
Dissecting germ cell heterogeneity at a single-cell resolution. **a** Subclustering germ cell populations from the tSNE projection of all cells from the gonads. All germ cells were subclustered using Seurat (left) and were then analyzed with PCA (right, color-coded with the cell identity). **b** tSNE projection of all germ cells. Cells from E11.5, E12.5 and E13.5 showed a closely clustering pattern while E14.5 showed a discrete clustering pattern (left). Reforming tSNE clustering illustrated 7 transcriptionally different sub-clusters (right). **c** Violin plot illustrating mitosis germ cell, early meiotic and late meiotic marker genes expression in each germ cell clusters. **d** Volcano plot demonstrating DEGs in different germ cell stages. Representative DEGs for each cluster were colour coded with corresponding cluster colours. Red dots represent significant DEGs while the grey dots represent non-significant DEGs. **e** Dot plot illustrating cluster-specific gene expression across all germ cell clusters. The dot size represents the percentage of cells expressing the indicated genes in each cluster and the dot colour intensity represents the average expression level of the indicated genes.

We next visualized a series of typical marker genes of germ cells to assign them within each cluster: *Dppa5a*, *Utf1*, *Pou5f1* and *Sox2* for mitotic (premeiotic) PGCs, *Stra8*, *Rec8*, *Dazl* and *Smc1b* for early meiotic germ cells, and *Dmrtc2*, *Tex12*, *Tex15* and *Taf7l* for the late meiotic germ cells (Figure 2c) (Handel & Schimenti, 2010). We found that mitotic PGCs mainly allocated in subcluster 2, 4 and 5, while early meiotic markers mainly expressed in clusters 0, 1 and 3, and late meiotic markers showed the highest expression in cluster 6. Combined with the developmental point for each cell clusters (Figure 2b, left panel), it’s plausible that cluster 3 marks the primary population of germ cells that have initiated meiosis, while cluster 6 marks the late stage of germ cells during meiosis prophase I. Since we have successfully identified four transcriptional distinct populations (cluster 3, 1, 0 and 6) that marks different stage of meiosis prophase I, we then wonder to investigate the sequential changes of all DEGs that drive the progression of meiosis by analysing the gene expression differences between different clusters. By analyzing the DEGs expressions among mitotic clusters (mitotic 0, cluster 2, 4, 5), meiotic I (cluster 3), meiotic II (cluster 1). meiotic III (cluster 0), and meiotic IV (cluster 6) that marks the progression of meiosis, we found that the earliest meiotic I stage significantly elevated classical meiosis “gatekeeper” genes *Stra8*, *Dazl*, *Dusp9* and *Smc1b* (encoding DNA recombination proteins) (Handel & Schimenti, 2010), while in the meiotic II stage, the expression of *Stra8*, *Smc1b*, and *Serf1* (encoding a protein with unknown function) was further elevated. Besides, synaptonemal complex protein family *Sycp1/3*, *Rec8* (a meiosis-specific component of the cohesins) (Xu et al., 2005) and *Taf7l* (well known for its role in regulating spermiogenesis) (Zhou et al., 2013) were upregulated at this stage. For the latter two stages, *Sycp3*, *Taf7l*, *Tex15* (reported to be expressed in male meiotic germ cells) (Tsukamoto et al., 2006, Yang et al., 2008), and *Dmrtc2* (doublesex and mab-3 related transcription factor-like family C2) (Handel & Schimenti, 2010) were continuously up-regulated. Noteworthy, *Prdm9* (a major determinant of meiotic recombination hotspots) (Baudat et al., 2010) was significantly up-regulated in the late stage of meiosis, while *Pou5f1* and *Rec8* significantly decreased at this stage.

Besides, we further characterized the germ cell stage-specific marker gene expression (Figure 2e). Similar to our analysis illustrated above, germ cells at meiotic I showed similar gene expression to mitotic stage germ cells except for high expression of *Dazl* and *Serf1*. As for germ cells at meiotic II and III, we observed elevated expression of meiosis marks such as *Stra8*, *Rec8* and *Tex12*, and decreased expression of mitotic cluster genes. For the meiotic IV stage germ cells, they specifically increased *Prdm9*, *Inca1*, *Sycp2*, *Syce3*, *Dmrtc2*, and significantly decreased mitotic and meiotic I, II, III stage marker genes. We also compared the cluster enriched GO terms and the Circos plot demonstrated that cluster 0 (meiotic III) and 6 (cluster IV) enriched many overlapped GO terms, including synaptonemal complex assembly and meiotic cell cycle (Supplementary Figure 3b, c). As for clusters 2, 4 and 5, they similarly enriched GO terms of mitotic cell cycle process and mitotic nuclear division (Supplementary Figure 3c), further confirmed their mitotic germ cell identity.

### Recapitulating gene regulatory networks (GRNs) underlying germ cell meiosis initiation

To recapitulate the gene regulatory relationships among the different germ cell clusters and infer the regulatory mechanisms underlying the germ cell status transition, we then used SCENIC, an algorithm developed to deduce GRNs and cellular status for scRNA data (Aibar et al., 2017). We, therefore, extracted the germ cell expression matrix from Seurat and imported them as the input matrix for SCENIC. By recognizing the co-expression modules (including transcriptional factors) and cis-regulatory motif analysis with each co-expression modules, we obtained a series of cell identity specific regulons together with their corresponding targets (Supplementary Table 2). Next, we used the AUCell algorithm to score the activity of each regulon in each cell according to the standard SCENIC pipeline and obtained the binary activity regulon matrix. To confirm our Seurat tSNE clustering based on the highly variable genes, we then reperformed cell clustering of germ cells based on the AUCell scored regulon activity as regulon activity was also cell identity specific as previously described (Aibar et al., 2017). Consistent with the Seurat tSNE algorithm, cell clustering using regulon activity also obviously distinguished mitotic, meiotic I, II, III and IV stage germ cells (Figure 3a), confirming the *bona fide* characterization of cell identify by using aforementioned tSNE clustering.

**Figure 3.**
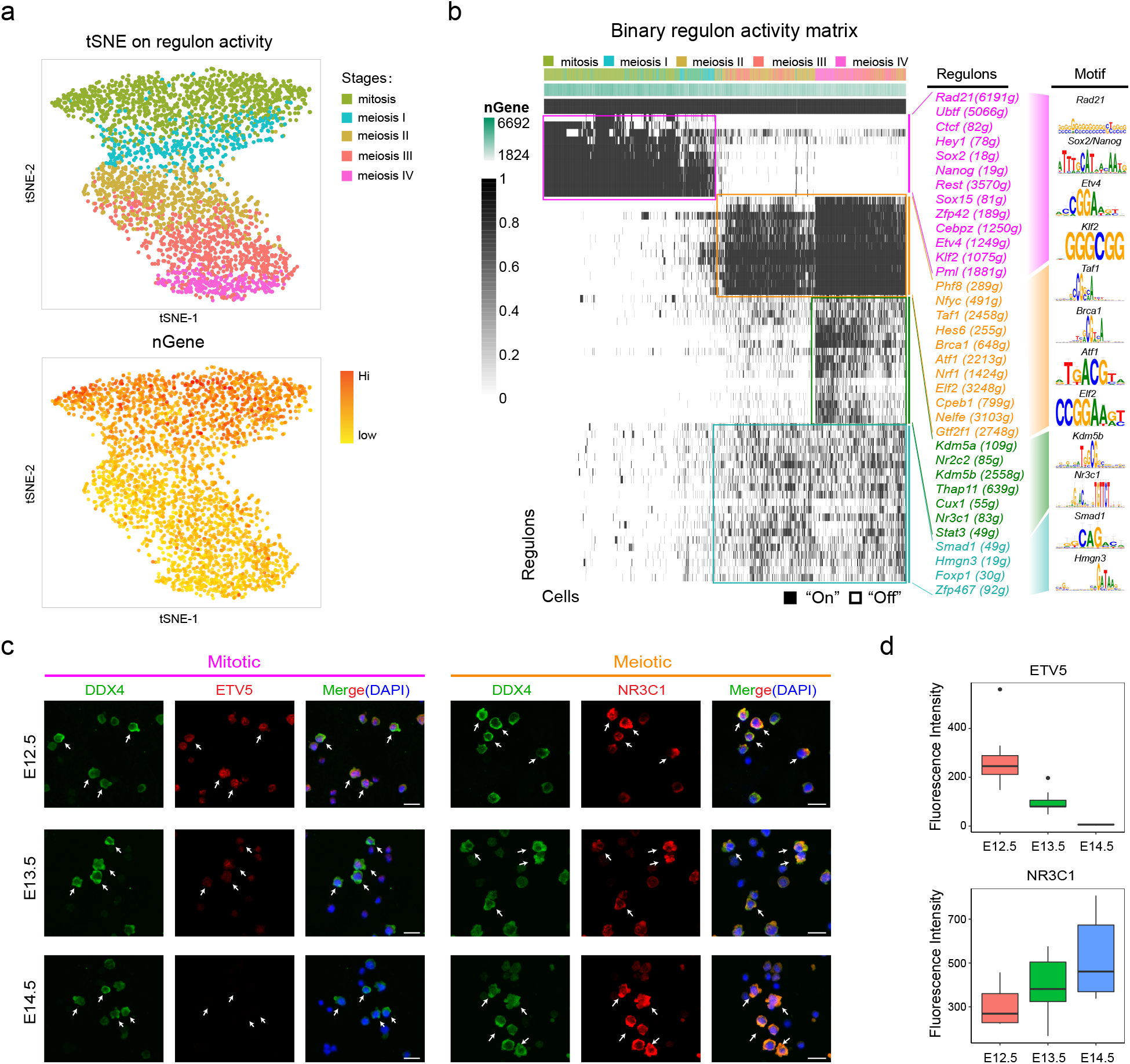
Single-cell regulatory network inference and clustering in the germ cell clusters. **a** tSNE projection of all germ cells based on the binary regulon activity. Top: Cells are labeled with their Seurat determined identity. Bottom: Gene density projected into the tSNE plot. The color density represents number of genes. **b** SCENIC binary regulon activity matrix showing the cell type-specific enrichment of regulons. Each column represents one single cell and each row represents one regulon. On” represents active regulons and “Off” represents inactive regulons. **c** Immunofluorescence analysis of mitotic regulon ETV5 and meiotic regulon NR3C1 in E12.5, E13.5 and E14.5 ovarian cells, germ cells were stained with DDX4 and white arrows indicates ETV5/NR3C1 and DDX4 double positive germ cells. Scale bars = 25 μm. **d** Fluorescence intensity analysis of ETV5 and NR3C1 expression at E12.5, E13.5 and E14.5.

Next, to recognize the master regulators within each cell population, we visualized regulon activity across all the germ cells based on the regulon scores calculated by AUCell. Noteworthy, the binary regulon activity heatmap indicated that mitotic germ cell clusters and meiotic I germ cell clusters predominantly showed high expression of regulons including *Sox2*, *Etv4* and *Nanog* regulons (Figure 3b), all of which were well defined regulators in the maintenance of pluripotency and self-renewal capacity of embryonic stem (ES) cells (Akagi et al., 2015). We also found that mitotic germ cells also enriched cell cycle-related regulon *Rad21and Rest* (Atienza et al., 2005, Zhang et al., 2017), which was consistent with their mitotic cellular status. Interestingly, some of these regulons gradually “turned off” in the meiotic I germ cells clusters, it’s therefore plausible that the down-regulation of mitotic specific expressed regulons (pluripotency and cell cycle-related) is pivotal for germ cells to initiate meiosis fate. Consistent with our analysis here, Yamaguchi *et al*., demonstrated that the NANOG protein was expressed in E11.5 and E12.5 germ cells while it was undetectable in E13.5 and E14.5 germ cells (Yamaguchi et al., 2005), also confirmed our analysis here. Besides, immunofluorescence analysis of ETV5 showed that ETV5 mainly expressed at E12.5, E13.5 (Figure 3c), while it was undetectable at E14.5 as revealed by fluorescence intensity analysis, further confirming our regulon analysis here (Figure 3d).

For meiotic germ cell clusters (meiotic II, III and IV stage), they similarly enriched a series of regulons, including *Phf8* (encoding a histone lysine demethylase) (Loenarz et al., 2010), *Taf1* (interact directly with TATA binding protein) (Metcalf & Wassarman, 2007), *Brca1* (highly expressed in pachytene and diplotene spermatocytes)(Zabludoff et al., 1996), *Elf2* (a rhombotin-2 binding ets transcription factor) (Wilkinson et al., 1997) etc. Noteworthy, a series regulon was gradually “turned on” meiotic IV stage germ cells, including *Kdm5a* (encoding an histone demethylase) (Varaljai et al., 2015), *Nr3c1* (encoding a glucocorticoid receptor with transcription or co-transcription factor functions) (Palma-Gudiel et al., 2015) and *Stat3* (critical for meiotic cell cycle) (Wang et al., 2018). Immunofluorescence of the meiotic regulon NR3C1 showed that NR3C1 was specifically expressed in germ cells, and that its expression increased with the progression of meiosis, thus deciphering potential roles during the progression of meiosis (Figure 3c and 3d).

### Pseudotime reconstruction of meiosis progression trajectory

According to the tSNE projection, we successfully identified mitotic PGCs, meiotic I, II, III and IV stage germ cells and delineated their detailed signature gene expression patterns during germ cell meiosis initiation (Figure 4a). Interestingly, the percentage of meiotic I stage germ cells significantly increased at E13.5, while the percentage of meiotic II, III and IV significantly increased at E14.5 (Figure 4b). To reconstruct the pseudotime trajectory of germ cells during meiotic progression, we used variable genes identified by Seurat as ordering genes (Supplementary Figure 5a) and performed pseudotime ordering of all germ cells using Monocle, a new and improved algorithm for classifying and counting cells, performing differential expression analysis between the subpopulations of cells (Qiu et al., 2017b). After construction of the germ cell lineage trajectory, we observed an inverted “U” structure with cells from E11.5 and E14.5 distributed at the two terminals (Figure 4c), which was also consistent with their relationship from the perspective of developmental timepoint. Besides, we also analyzed cell cluster distribution along pseudotime, and it was found that the precedence relationship identified according to their gene expression pattern also consistent with Monocle analysis here, with mitosis cell populations (cluster 2, 4, 5) distributed at the left and meiotic cell clusters (cluster 3, 1, 0, 6) distributed at the right (Figure 4d), further demonstrating the progression of meiosis.

**Figure 4.**
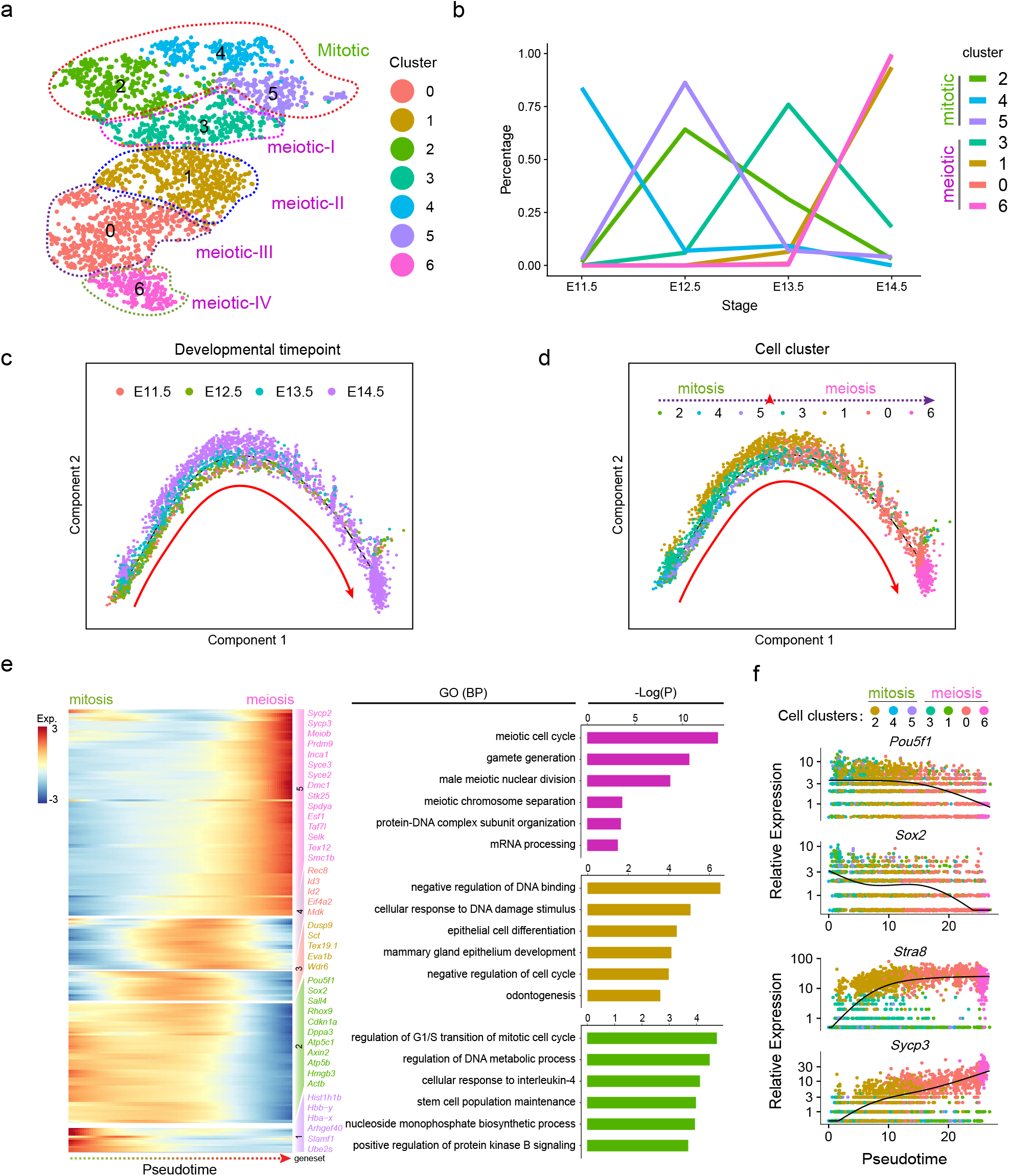
Pseudotime ordering of all germ cells along meiosis progression. **a** tSNE projection of all germ cell clusters. Germ cells were divided into mitotic, meiotic I, meiotic II, meiotic III, meiotic IV germ cell clusters. **b** Germ cell cluster dynamics during four different developmental time points. Clusters were color-labeled corresponding to the tSNE plot. **c** Pseudotime ordering of all germ cells colored by their cell identity. Each dot was colored according to their developmental time point. **d** Pseudotime ordering of all germ cells colored by their cell clusters. **e** Heatmap representing gene expression dynamics during pseudotime ordering of all germ cells and GO enrichment analysis of DEGs from different gene sets. **f** Pseudotime expression pattern of representative genes. Cells were coloured with their cluster information.

To gain in-depth insight into the gene expression dynamics during the initiation of meiosis, we analyzed gene expression dynamics along pseudotime and observed 5 distinct DEGs (*q-*val < 1e-4) sets according to *k*-means clustering (Figure 4e). We also performed GO enrichment to investigate the gene function categories of these DEGs and it was found that at the early stage (gene set2), germ cells significantly expressed genes enriched in GO terms of regulation of G1/S transition of mitotic cell cycle and regulation of DNA metabolic process, while in the middle stage (gene set 3), germ cells expressed genes mainly enriched in GO terms of negative regulation of DNA binding and cellular response to DNA damage stimulus. At the endpoint of pseudotime trajectory, germ cells expressed genes enriched in GO terms of meiotic cell cycle and gamete generation. Besides, we also evaluated pluripotent marker *Pou5f1*, *Sox2, Sall4, Dppa3, Eif4a1 and Rhox9* (Okashita et al., 2016) expression levels along pseudotime and found that *Pou5f1*, *Dppa3*, *Eif4a1* and *Rhox9* significantly decreased when germ cells initiated meiotic program while the expression of *Sox2* and *Sall4* continuously decreased along pseudotime (Figure 4f and Supplementary Figure 5b). For meiotic markers *Stra8*, *Sycp3*, *Prdm9*, *Sycp2*, *Taf7l* and *Tex12*, they both showed elevated levels along pseudotime, which was also consistent with our previous DEGs analysis. Noteworthy, the expression of *Prdm9* and *Sycp2* was elevated in the late stage of meiosis, thus emphasizing their roles in the late stage of meiotic progression. Taken together, our data here characterized detailed germ cell gene expression dynamics along meiosis progression and provided us in-depth insight into the molecular mechanisms underlying meiosis regulation.

### Dissecting heterogeneity and cellular fate decisions of the granulosa cell lineage

After the initial identification of the supporting and pregranulosa cell clusters, we then re-performed tSNE analysis on *Wnt4*, *Wnt6*, and *Fst* positive cell populations to preliminarily investigate the cellular heterogeneity of the identified supporting cell and pregranulosa cell clusters, as granulosa cells have been demonstrated to interact with an oocyte to promote oocyte development (Figure 5a and 5b) (Chang et al., 2016). tSNE projection resulted in 7 subclusters showing distinct developmental-dependent dynamics (Figure 5b lower panel and Supplementary Figure 6a), each characterized by a specific gene profile (Supplementary Figure 6b).

**Figure 5.**
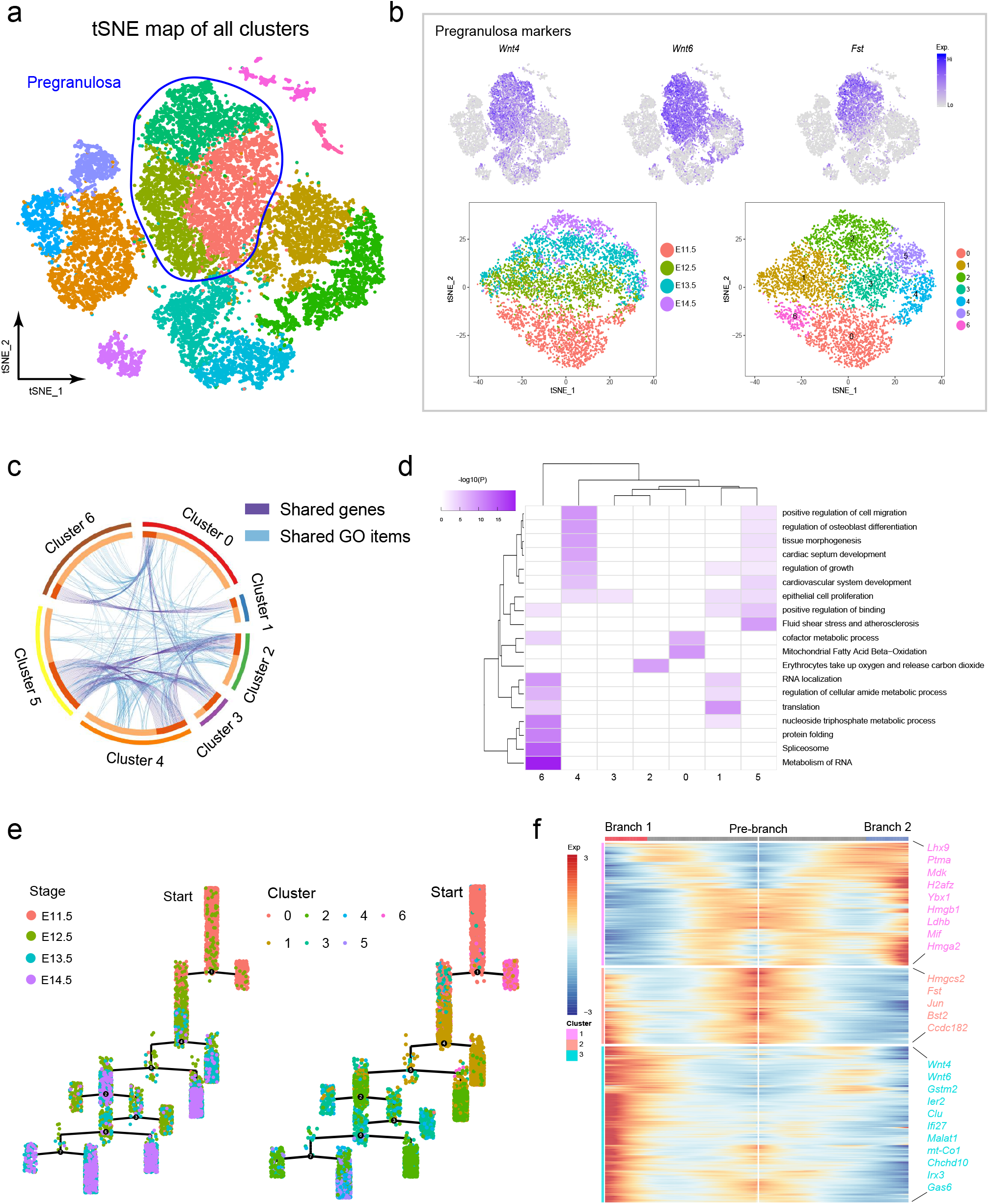
Dissecting granulosa lineage cellular heterogeneity along the progression of meiosis. **a** Pregranulosa cell population highlighted in the tSNE plot. **b** Top: Identification of pregranulosa cell markers *Wnt4*, *Wnt6* and *Fst* expression in the tSNE plot of all cell clusters. Bottom: tSNE projection of pregranulosa cell clusters. Cells are labelled with sample ID and cluster ID, respectively, and tSNE clustering reveals 6 subclusters in pregranulosa lineage cells. **c** Circos plot displaying shared DEGs and shared GO terms between different pregranulosa cell clusters. Shared DEGs were labeled with purple lines and shared GO terms were labeled with light blue lines. **d** Heatmap demonstrating the enrichment of GO terms in each pregranulosa cell cluster. Each row represents GO terms and each column represents cell clusters. **e** Pseudotime ordering of all pregranulosa cells by Monocle. The distance from a cell to the root corresponds to pseudo-time. **f** Heatmap representing gene expression dynamics during pseudotime ordering of pregranulosa cells. The branch point represents branch point 4 in Figure 5e.

The comparison of the shared DEGs and GO terms showed that subcluster 0 shared the GO terms of “cofactor metabolic process” with subcluster 6, while subcluster 2, 3, 4 and 5 shared more GO terms (Figure 5c and 5d). Combined with their cell label information, we found that subcluster 0, 1 and 6 were mainly composed of cells from E11.5 and E12.5, while subcluster 2, 3, and 5 were mainly composed of cells from E13.5 and E14.5 (Figure 5b), which likely delineates the differentiation of supporting cells into pregranulosa cells. We then reconstructed the pseudotime ordering of all granulosa cell lineage precursors and it was revealed that within the four developmental points, granulosa cell lineage precursors branched into multiple developmental branches (Figure 5e and Supplementary Figure 6c). This suggested a high heterogeneity of granulosa cell lineage precursors during meiotic initiation. To define more precisely the gene expression transition during the cell fate decision from supporting cells into pregranulosa cells, between E12.5 and E13.5, we performed DEG comparisons on the pseudotime ordering of these cells (Figure 5f). We observed three sets of DEGs and for the pre-branch lineage, DEGs enriched the GO terms of “organic acid catabolic process and small molecular biosynthetic process”, while branch 1 (characterized by elevated expression of *Wnt4* and *Wnt6*) and branch 2 (characterized by expression of *Lhx9* and *Ptma*) enriched the GO terms of “actin filament-based process, positive regulation of cell death” and “ribonucleoprotein complex biogenesis, translation”, respectively (Supplementary Figure 6d, 6e).

### Mesothelial and interstitial cell populations show two distinct cellular states

In our aforementioned tSNE projection of somatic cell clusters, it was observed that two mesothelial, two interstitial and one endothelial cell population(s) can be characterized (Figure 6a). Interestingly, the preliminary analysis indicated that the clusters within the interstitial and mesothelial cells expressed a similar level of *Bgn* and *Upk3b*, that we had used to mark interstitial and mesothelial, respectively (Figure 6b, top panel). However, the expression of cell cycle-related genes, such as *Cdk1*, *Ccna2* and *Cenpa*, showed specific expression only in interstitial cluster 2 and mesothelial cluster 4 (Figure 6b, lower panel). To gain further insight into the transcriptome differences between the cell clusters, we then compared the top100 expressed DEGs between the four clusters. The Venn diagram demonstrated that cluster 2 from interstitial cells and cluster 4 from mesothelial shared a very high percentage of overlapped DEGs (Figure 6c, left). Besides, protein to protein interaction network analysis suggested that the 41 co-expressed DEGs mainly enriched in “cell cycle-related network” (Figure 6c, right panel). GO analysis also indicated that the co-expressed DEGs enriched the GO terms of “cell cycle, cell division and regulation of mitotic cell cycle” (Figure 6d), suggesting that clusters 2 and 4 shared a similar cellular status. As for cluster 1 and cluster 3, GO enrichment analysis showed that cluster 1 enriched the GO terms of “cardiovascular system development, circulatory system development and tissue development” and cluster 3 enriched the GO terms of “organ development, system development, and anatomical structure development” (Supplementary Table 3). On the whole, these results suggest that during the early stages of ovary development interstitial and mesothelial cells possess two cellular states, one primed to differentiation and another to self-renew.

**Figure 6.**
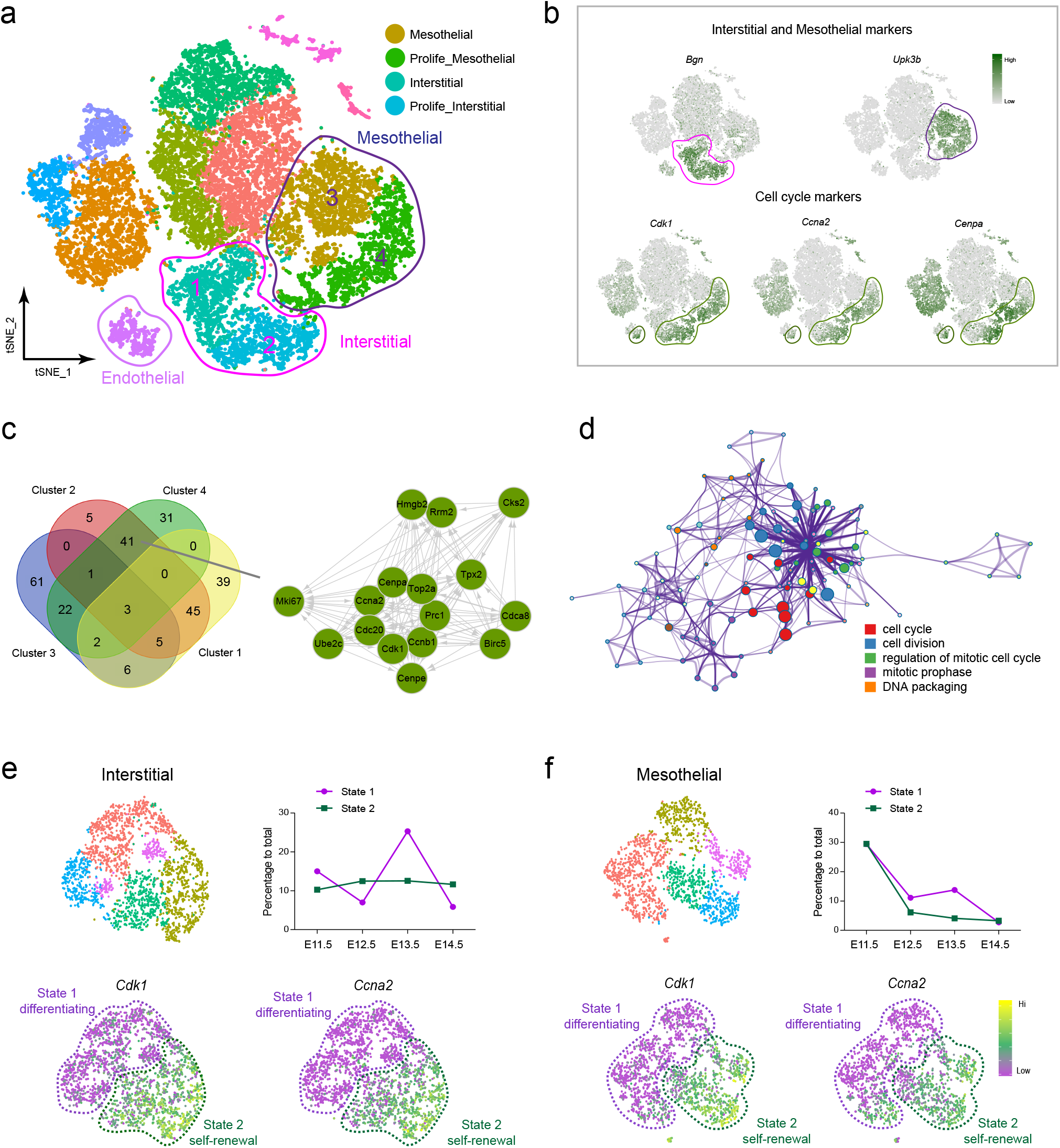
Dissecting interstitial and mesothelial heterogeneity by using scRNA seq. **a** Interstitial and mesothelial cell populations are highlighted in the tSNE plot. **b** Visualizing the expression of the interstitial marker (*Bgn*), mesothelial marker (*Upk3b*) and the cell cycle-related genes (*Cdk1*, *Ccna2* and *Cenpa*) in the tSNE plot. **c** Venn plot demonstrating the overlap of co-expressed DEGs between the interstitial and mesothelial subclusters. Cluster 2 and cluster 4 co-expressed DEGs were extracted to perform protein-protein network analysis using STRING database. **d** Enrichment network of co-expressed DEGs between cluster 2 and cluster 4. Each node represents one GO term and the node size represents the number of enriched genes. **e** Reperforming tSNE analysis on interstitial clusters and visualization of cell status dynamics and cell cycle-related marker genes (*Cdk1* and *Ccna2*). Interstitial clusters were extracted to reperform tSNE clustering. We termed subclusters expressing a high level of cell cycle-related genes as “State 2” and the remaining “State 1”. **f** Reperforming tSNE analysis on mesothelial clusters and visualization of cell status dynamics and cell cycle-related markers genes (*Cdk1* and *Ccna2*). Mesothelial clusters were extracted to reperform tSNE clustering. We termed subclusters expressing a high level of cell cycle-related genes as “State 2” and the remaining “State 1”.

**Figure 7.**
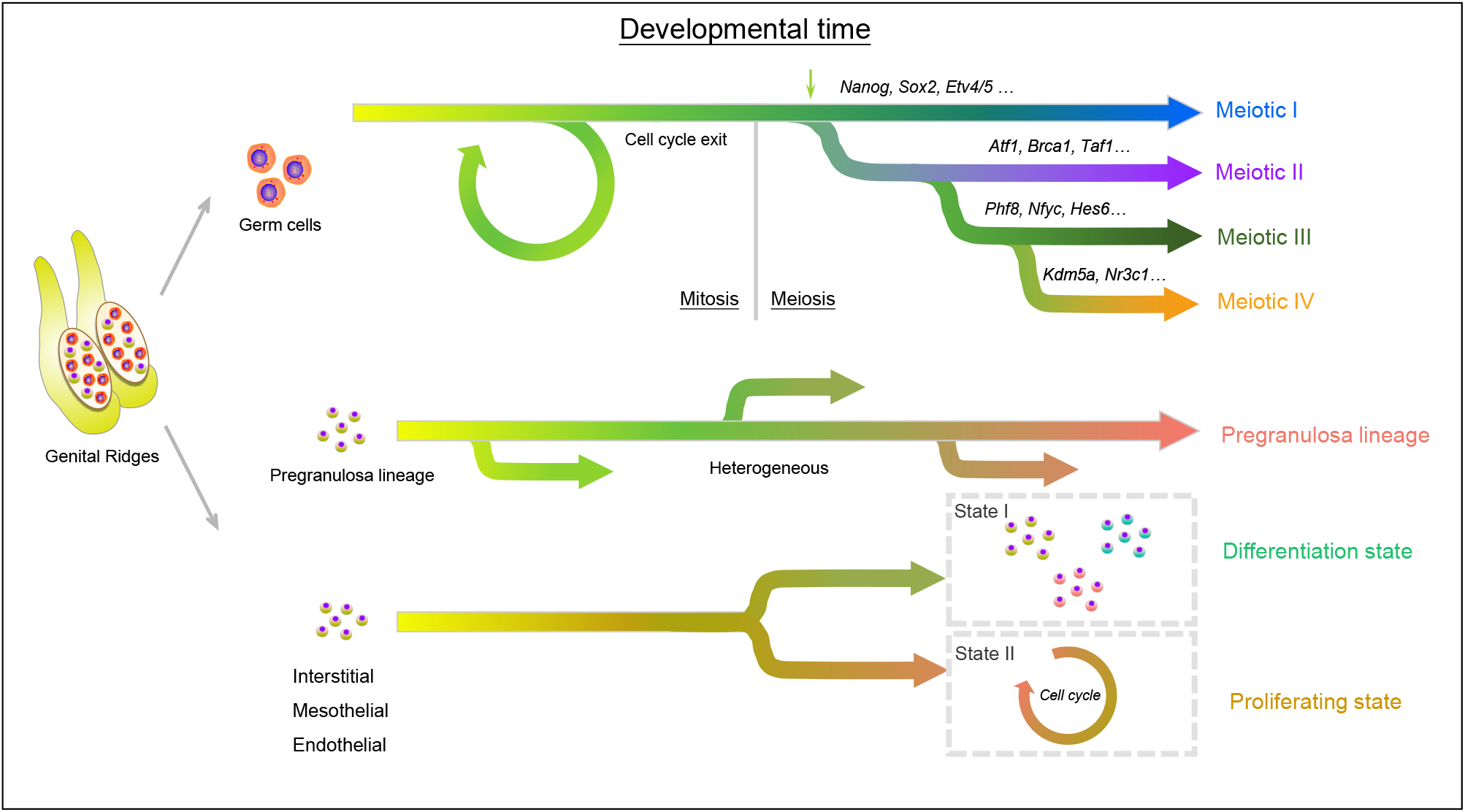
Model of the cellular dynamics in female gonads during the initiation of meiosis in germ cells. Germ cells are highly proliferative with a linage priority to meiosis initiation and express high levels of cell cycle related genes at E11.5 and E12.5. At around E13.5, a small population of germ cells initiate meiosis and this process is asynchronous in mice. Granulosa cell lineage cells are highly heterogeneous during this stage and the interstitial, mesothelial and endothelial cells reveal two cellular states, one showing “differentiating” characteristics, while the other displays “self-renewing” characteristics.

Finally, we analyzed the interstitial, mesothelial and endothelial cells at a higher resolution by extracting the three cell populations and re-performing tSNE clustering analysis. The tSNE projection revealed three subclusters for endothelial cells and five subclusters each for interstitial and mesothelial cells (Supplementary Figure 7a). We next compared the top five DEGs among the clusters and found that cluster 1, 2 in interstitial, cluster 2, 3, 4 in mesothelial cells and a portion of cluster 1, 2 in endothelial cells showed higher expression of cell cycle-related genes, such as *Top2a*, *Cenpa* and *Cdk1* (Supplementary Figure 7b, 7c). This illustrated that interstitial, mesothelial and also endothelial populations possess two cellular states. Furthermore, it was found that the percentage of interstitial cells with “differentiating” status was higher at E13.5 while the “self-renewal” state remained quite constant (Figure 6e). On the other hand, in the mesothelial cells the percentage of both states progressively decreased alongside developmental time (Figure 6f).

## DISCUSSION

The scRNA-seq technology facilitates the identification of new cell types and gene regulatory networks as well as allowing dissection of the kinetics and patterns of allele-specific gene expression (Bacher & Kendziorski, 2016, Liu & Trapnell, 2016). In such studies, individual cells are executing gene expression programs in an unsynchronized manner. Single cell gene expression studies enable profiling of transcriptional regulation during complex biological processes and within highly heterogeneous cell populations. These studies allow the discovery of genes that identify certain subtypes of cells, or that mark an intermediate status during a biological process (Rheaume et al., 2018).

In the present paper, we used such an approach for the first time in order to obtain new insights into molecular events underlying meiosis initiation and progression into prophase I in female mouse germ cells. We used an experimental model to characterize gene expression profiles of the heterogonous somatic cell populations present within the sex differentiating ovaries. To validate such analyses and separate biological variability from the possible technical noise that might affect scRNA-seq protocols, we employed various algorithms and bioinformatics analyses. The soundness of our analyses was confirmed, first of all, by utilizing our acquired understanding of the periods of PGC proliferation (E11.5-E13.5), followed by the initiation and progression of meiosis (E13.5-E14.5) in the female mouse fetus. This allowed using the expression of marker genes by PGCs before (*Pou5f1*, *Sox2* and S*all4*), and at meiotic initiation (*Dazl*, *Stra8, Smc1b* and *Rec8*), and later by oocytes during the first stages of meiotic prophase I (*Dmrtc2, Tex12, Tex15* and *Taf7l*).

The tSNE algorithm projection of the transcriptome data allowed clustering of four distinct populations of premeiotic PGCs and four germ cell populations in meiotic prophase I and their dynamics throughout the developmental period studied (Butler et al., 2018). Several new genes, highly expressed by each of these populations were also found. In total, 634, 654 and 1,189 DEGs were identified in premeiotic, early and late meiotic germ cells, respectively. By focusing on the germ cell populations using tSNE, we observed 7 germ cell clusters along meiosis progression. Noteworthy, we characterized 4 transcriptionally distinct meiotic germ cell clusters during meiosis prophase I. By analyzing their DEGs expression along meiosis progression, we delineated detailed gene expression landscape during meiosis progression in mice. For the meiotic initiation, our analysis demonstrated that *Stra8*, *Dazl*, *Dusp9*, *Hells* and *Serf1* were significantly up-regulated DEGs in meiotic I stage germ cells. For the late stage of meiosis, we observed DEGs such as *Pigp*, *Sycp3*, *Prdm9*, *Taf7l* and *Syce3*. Noteworthy, except for the classical *Stra8*, *Dazl*, *Sycp3* and *Prdm9*, the roles of *Dusp9*, *Hells*, *Serf1*, *Pigp* and *Taf7l* in meiosis progression remains little known and future studies aimed to unravel the role that some of these genes play during meiosis should provide critical information about early gametogenesis in mammals.

We also identified master regulons of germ cells which may drive the meiotic progression in mitotic, meiotic I, II, III and IV stage germ cells. For example, regulons such as *Etv4*, *Sox2* and *Nanog*, involved in the maintenance of pluripotency and self-renewal capacity of ES cells were predominantly expressed in mitotic PGCs, confirming their similarities with ES cells (Kalkan et al., 2019, Nicholas et al., 2009). While it could be expected that late meiotic germ cells, which involved extensive chromatin rearrangement, express the regulon *Hmgn3* encoding a nucleosome-binding protein thought to modulate the compactness of the chromatin fiber. The meaning of the expression of *Nr3c1*, encoding a glucocorticoid receptor with transcription or co-transcription factor functions, is intriguing but remained unexplained. Similarly, the implications of the prevalent expression in late meiotic germ cells of the regulons *Phf8*, a histone lysine demethylase, *Nfyc* a transcription encodes one subunit of a trimeric complex, *Kdm5a*, a histone demethylase and *Nelfe*, encodes a protein represses RNA polymerase II transcript elongation, remain to be investigated. Collectively, these analyses here reveal the unique GRNs and master regulons in germ cells at different cellular stages during meiosis progression.

Our research also provides detailed transcriptome profiles of the four ovarian somatic cell lineages differentiating in the embryonic ovary during the initiation and progression of meiosis, namely pregranulosa, interstitial, mesothelial and endothelial cells. For pregranulosa cells, the package Monocle revealed multiple differencing branches supporting a high initial heterogeneity of cells in this lineage. In addition, interstitial cells and mesothelial cells showed two distinct cellular states that according to their transcriptome profile, we defined as “self-renewing” and “differentiating”.

In conclusion, the present data represents a new important resource for deciphering the molecular pathways driving meiosis initiation and progression in female germ cells and ovarian somatic cells. Thereby, improving our understanding of the embryonic processes involved during gonadal development in female mammals. These results, prospectively, provide valuable information about the aetiology of human reproductive defects arising from dysregulation during early gametogenesis.

## MATERIALS AND METHODS

### Animals

All mice used in this study were C57/BL6 mice purchased from Beijing Vital River Laboratory Animal Technology Co., Ltd. Experimental procedures involved in this study were approved by the Animal Care and Use Committee of Qingdao Agricultural University. Briefly, all C57/BL6 mice were housed in a light and temperature-controlled room (light: 12 h dark and 12 h light cycles; temperature: 24 ± 0.5 °C) with ad libitum access to food and water. Then, 6-week old female mice were mated with 8-week old male mice (3:1) overnight and the vaginal plug was checked the next morning. Mice with a vaginal plug were considered 0.5 days post coitum (dpc).

### *In vitro* isolation of genital ridges and sex genotyping

Pregnant mice were sacrificed by cervical dislocation and the genital ridges of the fetus were isolated using a pair of precise forceps as previously described (Morohaku et al., 2016). For the characterization of E11.5 fetal ovarian tissues, the *Sry* and *Ube1* genes was used for sexing. The following primers were used according to the previously described procedure: *Sry*: F: 5’-CTG TGT AGG ATC TTC AAT CTC T-3’; R: 5’-GTG GTG AGA GGC ACA AGT TGG C-3’ and *Ube1*: F: 5’-TGG TCT GGA CCC AAA CGC TGT CCA CA-3’; R: 5’-GGC AGC AGC CAT CAC ATA ATC CAG ATG-3’ (Chuma & Nakatsuji, 2001). Briefly, a small piece of skin tissues of each fetus was isolated and was boiled in water for 10 min, then the tissues were directly used as PCR templates and PCR was performed using 2×EasyTag PCR SuperMix (Transgene, Beijing, China, AT311-03) according to the manufacturer’s instructions. The obtained PCR products were then electrophoresed on a 2 % agarose gel (TSINGKE, Beijing, China, R9012LE) at 100 V for 30 min (*Sry*) or 1h (*Ube1*). *Gapdh* (F primer: 5’-AGG TCG GTG TGA ACG GAT TTG-3’; R primer: 5’-TGT AGA CCA TGT AGT TGA GGT CA −3’) was used as a loading control. For sex determination of E12.5, E13.5 and E14.5 genital ridges, the sex was morphologically distinguishable according to the formation of testis cords in the male gonads (Li et al., 2014).

### Synaptonemal complex spreading assay

Immunofluorescence staining of synaptonemal complexes was used for determining the meiotic progression of germ cells at different stages as we previously described (Feng et al., 2014, Ge et al., 2015b). Briefly, the isolated genital ridges were incubated with hypotonic solution (30 mM Tris, 50 mM sucrose, 17 mM citric acid, 5 mM EDTA, 2.5 mM DL-Dithiothreitol and 1 mM Phenylmethanesulfonyl fluoride in water) for 30 min at room temperature, after that, the genital ridges were transferred into 4 % paraformaldehyde solution (Sorlabio, Beijing, China, P1110) and the ovarian tissues were mechanically separated. The suspended ovarian cells were then spread onto glass slides overnight. The next morning, the slides were washed with 0.04 % Photo-Flo 200 (Kodak, Rochester, NY, USA, 146 4502) and then blocked with PBS supplemented with 1 % goat serum (BOSTER, Wuhan, China, AR0009) and 0.05M Tris-HCl. After blocking, the first antibody (SCP3, Novus Littleton, CO, USA, NB300-232; γH2AX, Abcam, Shanghai, China, ab26350) was then added and the slides were incubated at 37 °C for 8 h. After three times wash to remove unconjugated antibodies, the secondary antibodies (Donkey anti-Rabbit IgG H&L Alexa Fluor^®^ 555, Abcam, ab150074; Goat anti-Mouse IgG H&L Alexa Fluor^®^ 488, Abcam, ab150113) were then incubated at 37 °C for 2 h and the slides were mounted with Vectashield mounting media (Vector Laboratories, Burlingame, CA, USA, H-1000). All pictures were taken using a Leica Laser Scanning Confocal Microscope imaging system (Leica TCS SP5 II, Wetzlar, Germany).

### Immunofluorescence analysis and fluorescence intensity analysis

The isolated single cell pellets were first fixed with 4 % paraformaldehyde at 4 °C for 30 min, then, the cell pellets were plated on 3-Aminopropyl-Triethoxysilane (APES, ZSbio, Beijing, China, ZLI-9001) treated slides. Permeabilization was performed with PBST solution consisting of PBS supplemented with 0.5 % Triton X-100 (Solarbio, Beijing, China, T8200) for 10 min at room temperature. After permeabilization, slides were blocked with PBST supplemented with 10 % goat blocking serum (BOSTER, AR0009) for 45 min at room temperature. Primary antibodies (DDX4/MVH, Abcam, ab27591; UTF1, Abcam, ab24273; ETV5, Abcam, ab102010; NR3C1, Abcam, ab2768) were diluted in blocking buffer and were incubated with the slides at 4 °C overnight. The next morning, the slides were washed three times with PBS supplemented with 1 % BSA, and then the secondary antibodies were added and were incubated with the slides at 37 °C for 2 h. Finally, the slides were mounted with Vectashield mounting media and pictures were taken using a Leica Laser Scanning Confocal Microscope imaging system. The fluorescence intensity analysis was analyzed with ImageJ software (v 1.48, National Institutes of Health, Bethesda, MD, USA) according to the manufacturer’s instructions.

### Single cell library preparation and sequencing

Single cell library was prepared using the 10x Genomics Chromium Single Cell 3′ Library & Gel Bead Kit v2 (10×Genomics, Pleasanton, CA, USA, 120237) according to the manufacturer’s instructions. Briefly, to obtain the desired number of cells from genital ridges, about 8 - 10 female fetus were prepared for each group. The genital ridges were then mixed and dissociated with a 0.25 % trypsin-EDTA solution for 3 min at 37 °C. After trypsinization, the cell suspension was filtered with a 40 μm cell strainer (BD Biosciences, San Jose, CA, USA, 352340) and was washed three times with PBS solution supplemented with 0.04% bovine serum albumin (BSA, Sigma, St. Louis, MO, USA, A1933). To determine whether the cells obtained were eligible (cell viability > 80 %, cell concentrations = 1000 cells/μl) for downstream analysis, the cell viability was evaluated using trypan blue staining with a haemocytometer (Bio-Rad, Hercules, CA, USA, TC20) and the cell concentration was adjusted to 1000 cells/μl before loading to the single cell chip. The Gel-Bead in Emulsions (GEMs) were then generated with Chromium 10×Single Cell System (10×Genomics). To barcode cDNA in each cell, the cells were then lysed and followed by a reverse transcription procedure. After that, cDNA recovery was performed using DynaBeads MyOne Silane Beads (Invitrogen, Carlsbad, CA, USA, 37002D) according to the manufacturer’s instructions. cDNA libraries were then prepared using 10×Genomics Chromium Single Cell 3′ Library & Gel Bead Kit v2 following the manufacturer’s guide and sequencing was performed with an Illumina HiSeq X Ten sequencer (Illumina, San Diego, CA, USA) with pair-end 150 bp (PE150) reads.

### Single sample analysis and aggregation

CellRanger v2.2.0 software (https://www.10xgenomics.com/) was used to analyze the obtained datasets with ‘--force-cells = 5000’ argument to obtain the same number of cells for downstream analysis. The 10×Genomics pre-built mouse genome for mm10-3.0.0 (https://support.10xgenomics.com/single-cell-gene-expression/software/downloads/latest) was used as the reference genome. After the CellRanger pipeline, the gene-barcode matrices were then analyzed with Seurat single cell RNA seq analysis R package (v3.0) (Butler et al., 2018). The analysis procedure was performed according to the package’s user guide with little modifications. Briefly, cells with minimal genes less than 200 and genes expressed in less than 3 cells were removed to keep high-quality datasets for downstream analysis. After normalization, the four datasets were then merged together with Seurat RunMultiCCA function. To characterize cell clusters, they were visualized with Seurat RunTSNE function based on the t-distributed Stochastic Neighbor Embedding (tSNE) algorithm with default settings and cell clusters were calculated with FindClusters function at a resolution of 0.6. To characterize cell cluster markers the Seurat FindAllMarkers function was used.

### Subclustering, Gene Ontology and protein - protein network enrichment analysis

After the characterization of all cell clusters in genital ridges, cell clusters were further divided into different groups according to their cell identity. To extract the same type of cells for downstream analysis, we used SubsetData function implemented in Seurat package to extract germ cell and somatic cell subclusters. The extracted subclusters were then reanalyzed with the single Seurat analysis procedure to gain further insight into the subcluster information of particular cell types. After clustering and subclustering, cluster-specific markers were obtained with the FindAllMarkers function, to identify the intergenic relationship within clusters, we selected the cluster-specific top 500 differentially expressed genes based on *p*-value and performed Gene Ontology (GO) analysis using Metascape (http://metascape.org). To infer protein - protein network from different gene sets, STRING database (https://string-db.org/) was used to infer protein - protein interaction network and the network was further visualized with Cytoscape software (v3.7.0, https://cytoscape.org/).

### Single-cell pseudo-time trajectory analysis

Single-cell pseudo-time trajectory analysis was performed using R package Monocle 2 (v2.8.0) according to the online tutorials (http://cole-trapnell-lab.github.io/monocle-release/tutorials/) (Qiu et al., 2017a, Trapnell et al., 2014). Briefly, Monocle object was directly constructed using Monocle implemented newCellDataSet function from Seurat object with lowerDetectionLimit = 0.5 and we used the Seurat determined variable genes as highly variable genes for ordering. Dimensionality was reduced using the DDRTree method with regression based on the number of UMIs. The root state was chosen according to their Seurat cell identity information and branch-specific gene expression was calculated using Monocle implemented BEAM function and branched heatmap was further visualized by “plot_genes_branched_heatmap” function.

### Single cell regulatory network inference and clustering

To reveal gene regulatory networks during germ cell meiosis initiation, we performed regulatory network inference and clustering based on SCENIC, a modified method for inferring gene regulatory networks from single cell RNA seq data (Aibar et al., 2017). The SCENIC analysis was performed according to the SCENIC online tutorial (https://github.com/aertslab/SCENIC). Firstly, we extracted the single-cell RNA-seq expression matrix from Seurat, in which each column represents a cell ID and each row represents a gene, then we used geneFiltering function to remove genes with UMI counts across all samples less than 80.12 and expressed in less than 1 % of cells. After that, we used GENIE3 to infer co-expression matrix which contains potential regulators. To identify potential direct-binding targets, RcisTarget was then used based on DNA-motif analysis and we used databases (mm10) that score the motifs in the promoter of the genes (up to 500 bp upstream the TSS), and in the 10 kb around the TSS (+/-10 kb). Last, we used the AUCell algorithm to calculate regulon activity in each cell and convert the network activity into ON/OFF (binary activity matrix) with default settings.

## DATA AVAILABILITY

The sequencing raw data have been deposited in NCBI’s Gene Expression Omnibus (GEO) under accession number: GSE128553.

## SUPPLEMENTARY DATA

Supplementary Data are available online.

## ACKNOWLEDGEMENT

We would like to thank all members of Institute of Reproductive Sciences, Qingdao Agricultural University for their kind help during preparing single cell samples and suggestions for preparing the manuscript.

## Supplementary Figure legends

**Supplementary Figure 1** Quality control of the data set and representative cell marker gene expression analysis. **a** The summary metrics describe sequencing quality and various characteristics of the detected cells. **b** Violin plot demonstrating the number of genes (nGene), unique molecular identifier (nUMI) and percentage of mitochondria genes (percent.mito) in the different data sets. **c** tSNE plot of all single-cells coloured by sample information. **d** Marker gene expression projected into the tSNE plot. Mesothelial: *Lhx9*, *Upk3b*; Interstitial: *Col1a2*, *Bgn* Endothelial: *Pecam1*, *Kdr*; Pluripotent genes: *Utf1*, *Sall4*, *Pou5f1*, *Sox2*; Meiotic genes: *Sycp1*, *Rec8*, *Tex14*, *Mael*; Pregranulosa genes: *Wnt4*, *Wnt6*; Erythroid genes: *Alas2*, *Alad*; Immune genes: *Cd52*, *Car2*.

**Supplementary Figure 2** Interpreting cellular diversity in the gonads. **a** The cell population dynamics during four different developmental time-points. The cells were colored coded with the same color as in Figure 1d. **b** Heatmap of top 10 expressed DEGs in each cluster. Each color bar in the column represents one cell cluster, and each row represents one gene. **c** Dot plot illustrating marker gene expression across all cell types. The dot size represents the percentage of cells expressing the indicated genes in each cluster and the dot colour intensity represents the average expression level of the indicated genes. **d** Immunofluorescence analysis of the mitotic GC marker UTF1 in E12.5, E13.5 and E14.5 ovarian cells. Scale bars = 25 μm. **e** GO enrichment of DEGs in germ cell clusters 1, 8, and 9. **f** GO enrichment of DEGs in granulosa precursor clusters. **g** GO enrichment of DEGs in interstitial cell clusters. **h** GO enrichment of DEGs in mesothelial cell clusters. **i** GO enrichment of DEGs in endothelial cell clusters.

**Supplementary Figure 3** Dissecting germ cell subclusters. **a** Heatmap illustrating germ cell subcluster top 5 expressed genes. **b** Circos plot displaying shared DEGs and shared GO terms between different germ cell clusters. Shared DEGs were labeled with purple lines and shared GO terms were labeled with light blue lines. **c** Heatmap demonstrating the enrichment of GO terms in each germ cell cluster. Each row represents GO terms and each column represents a cell cluster.

**Supplementary Figure 4** SCENIC binary regulon activity matrix displaying all enriched regulons across different germ cell clusters. Each column represents one single cell and each row represents one regulon.

**Supplementary Figure 5** Pseudotime ordering of germ cells. **a** Visualization of variable genes used for Pseudotime ordering. **b** Expression profiles of representative marker genes identified by Monocle. The x-axis represents pseudotime.

**Supplementary Figure 6** Interpreting granulosa cell lineage cell fate using Monocle. **a** Pregranulosa cell lineage subclusters dynamics along developmental time point. **b** Heatmap of top 5 expressed DEGs in each cluster. Each column represents one cell cluster, and each row represents one gene. **c** Pseudotime ordering of all granulosa cell lineage clusters colored by cell identity, cell clusters, and cellular states. **d** GO enrichment of DEGs in different gene sets in Figure 5f. **e** Expression profile representative genes corresponding to each gene sets in Figure 5f.

**Supplementary Figure 7** Interpreting endothelial, interstitial and mesothelial cellular heterogeneity. **a** tSNE clustering of endothelial, interstitial, and mesothelial cell clusters. Cells were labeled with sample identity. **b** Heatmap of top 5 expressed DEGs in each cluster corresponding to endothelial, interstitial and mesothelial cells. Each column represents one cell cluster, and each row represents one gene. **c** Expression of cell cycle-related marker genes in tSNE projection of endothelial, interstitial and mesothelial cells.

## Supplementary Tables

**Supplementary Table 1.** List of DEGs in mitotic PGCs, early meiotic germ cells, and late meiotic germ cells.

**Supplementary Table 2.** Motif Enrichment analysis of significantly enriched regulons and potential targets.

**Supplementary Table 3.** Cluster-specific enriched genes in all cell clusters identified in gonads and the corresponding GO terms.

