## Supplemental Figures 1-7 for "Dissecting the initiation of female meiosis in the mouse at single-cell resolution"

### Supplementary Figure 1

a

| Sample info. | E11.5 | E12.5 | E13.5 | E14.5 |
|---|---|---|---|---|
| Estimated Number of Cells | 5,000 | 5,000 | 5,000 | 5,000 |
| Valid Barcodes | 97.4% | 97.2% | 97.3% | 97.0% |
| Mean Reads per Cell | 123,807 | 115,609 | 111,367 | 107,550 |
| Median Genes per Cell | 3,613 | 3,409 | 3,097 | 3,461 |
| Total Genes Detected | 19,932 | 19,862 | 19,959 | 20,033 |
| Reads Mapped to Genome | 94.20% | 94.60% | 93.80% | 94.80% |
| Reads Mapped Confidently to Transcriptome | 71.50% | 67.40% | 67.10% | 68.70% |

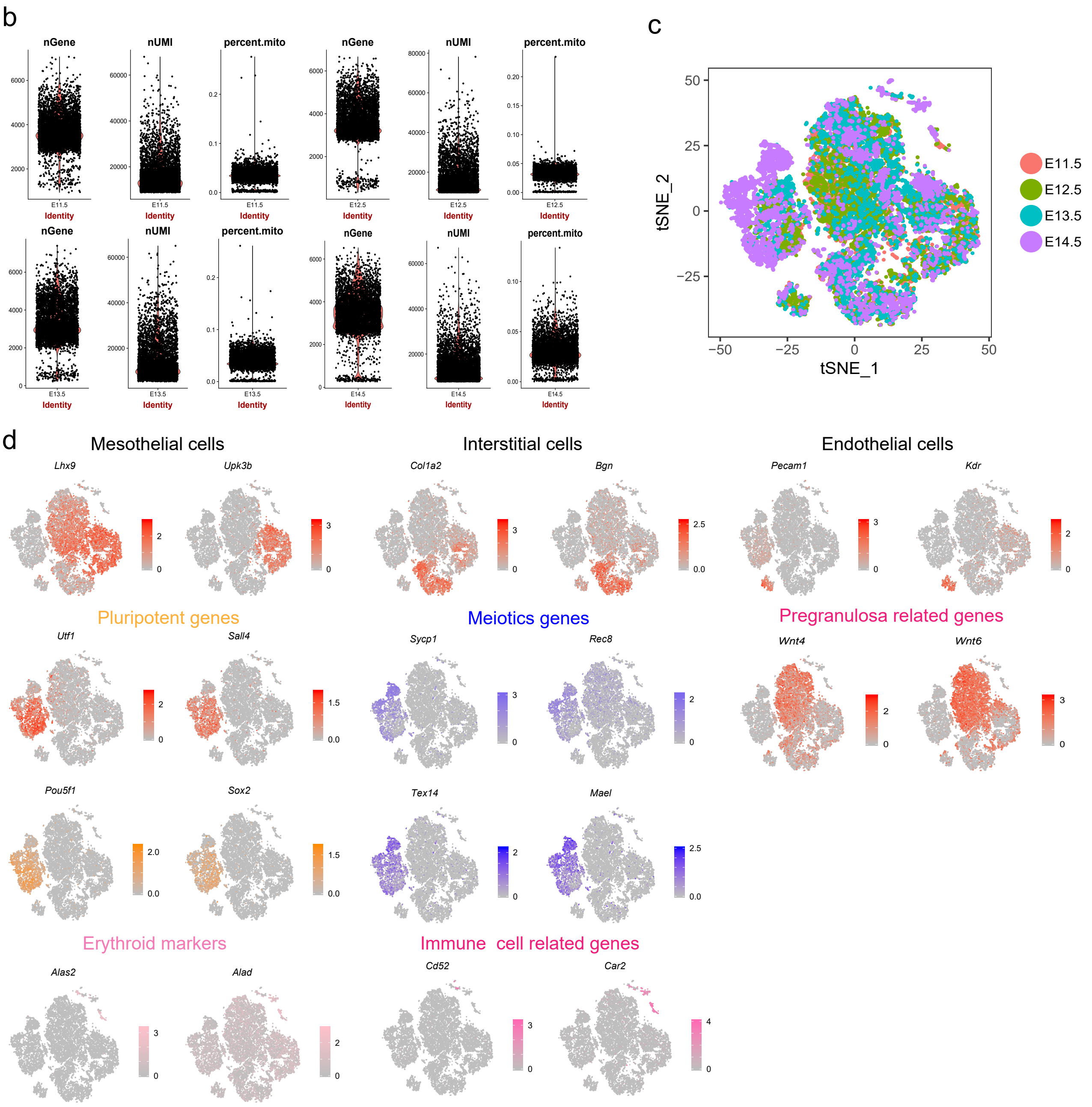

Supplementary Figure 2

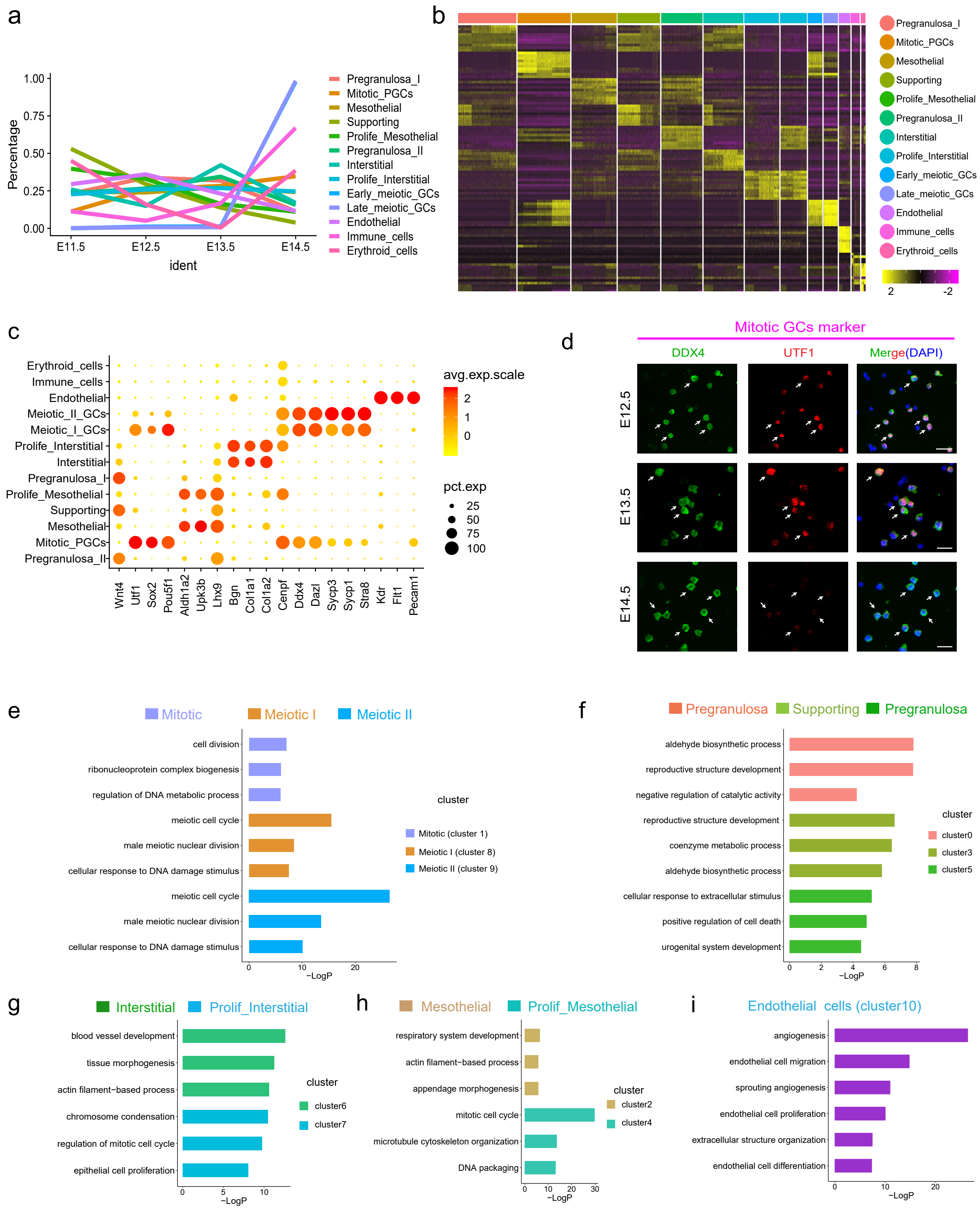

### Supplementary Figure 3

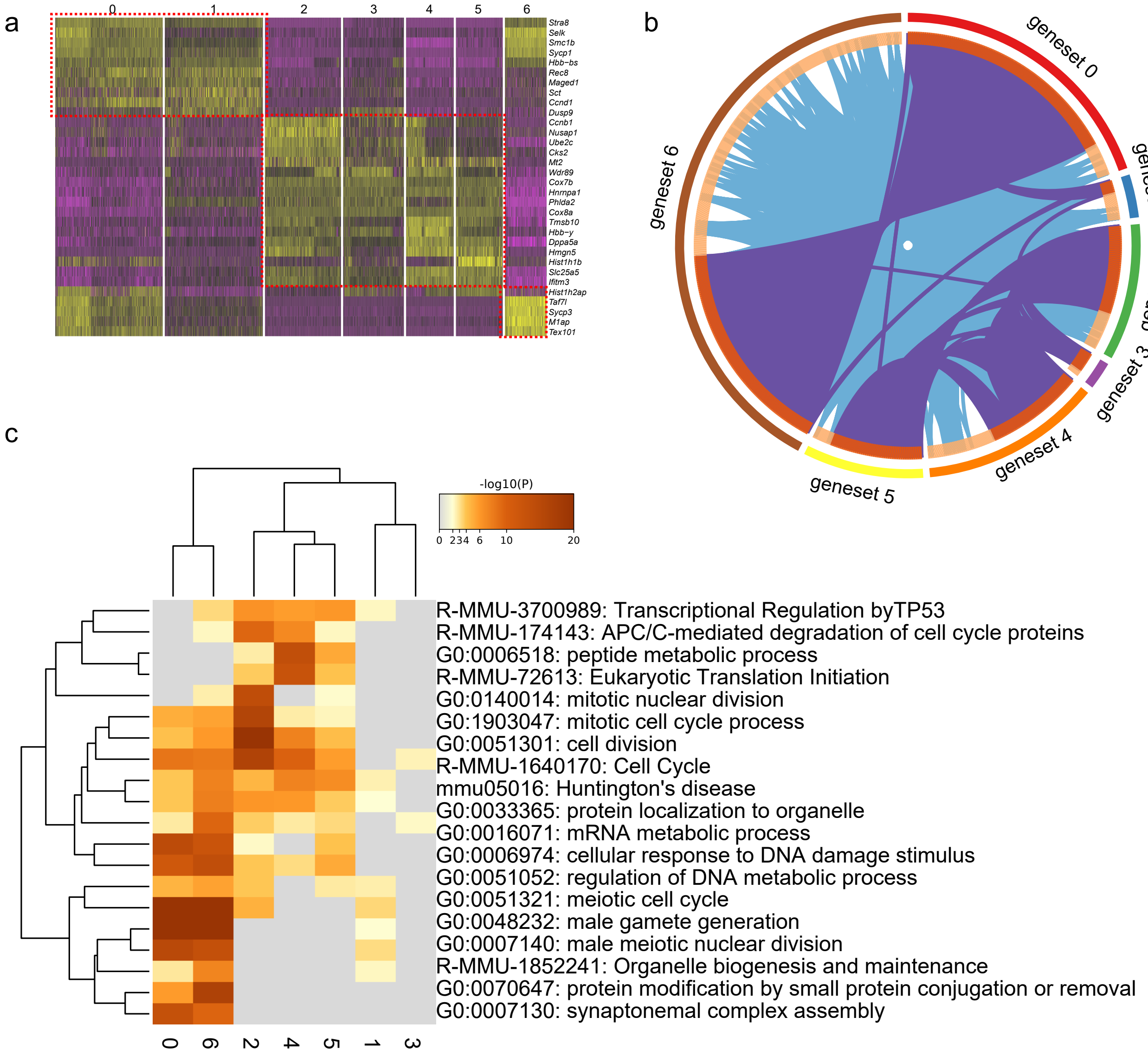

Supplementary Figure 4

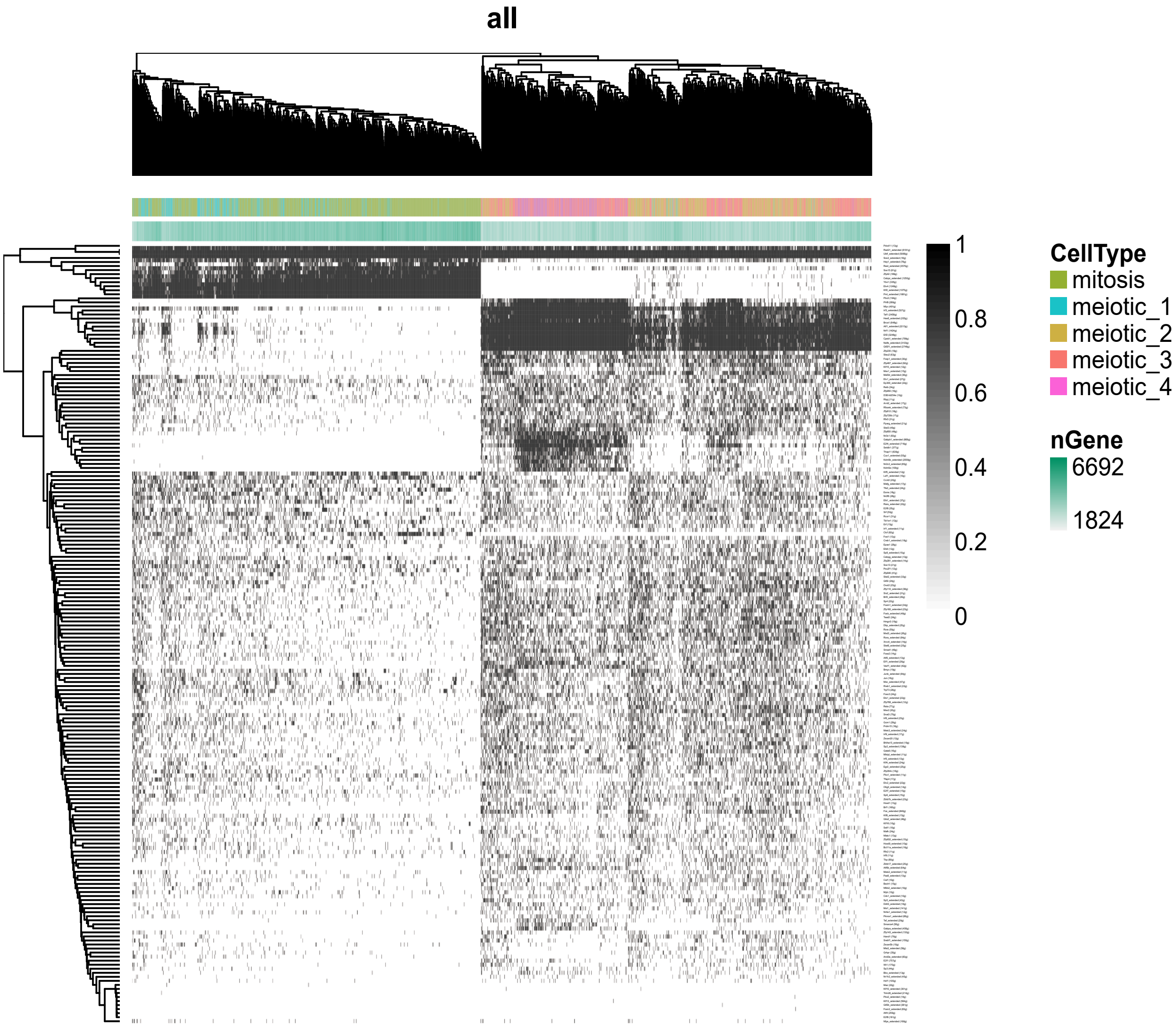

Supplementary Figure 4

a

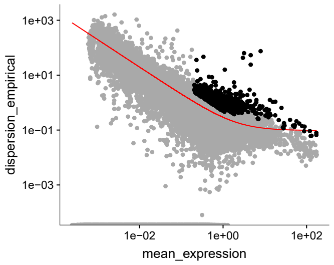


b

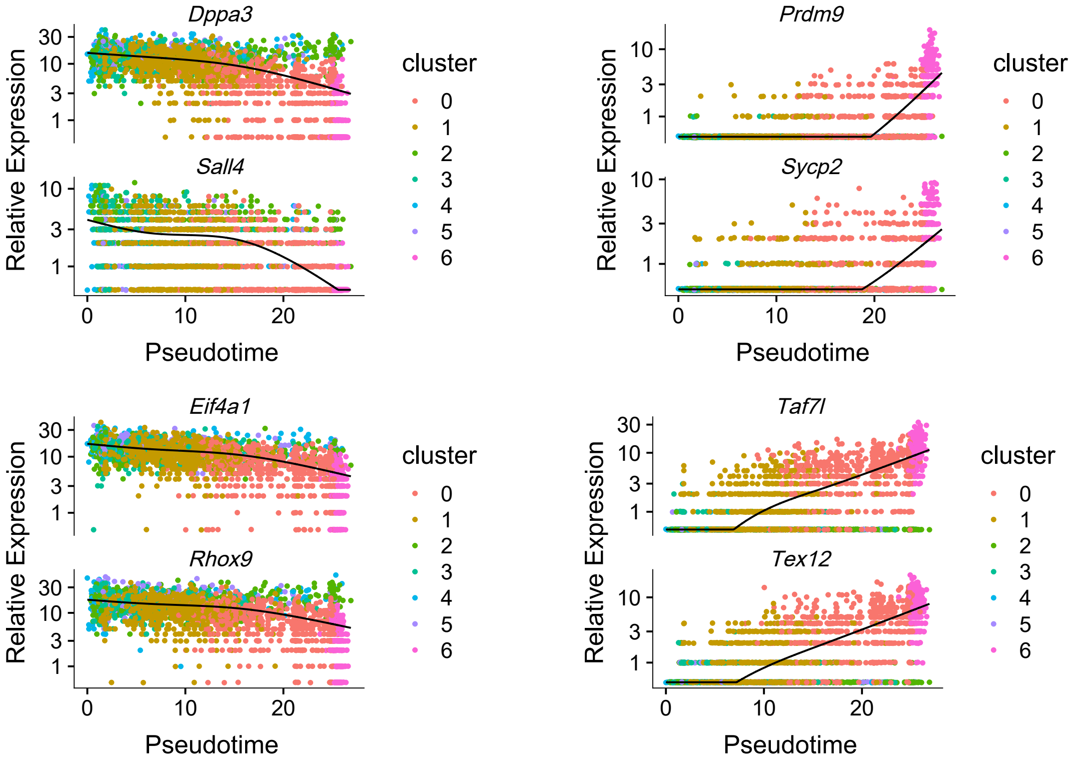

### Supplementary Figure 6

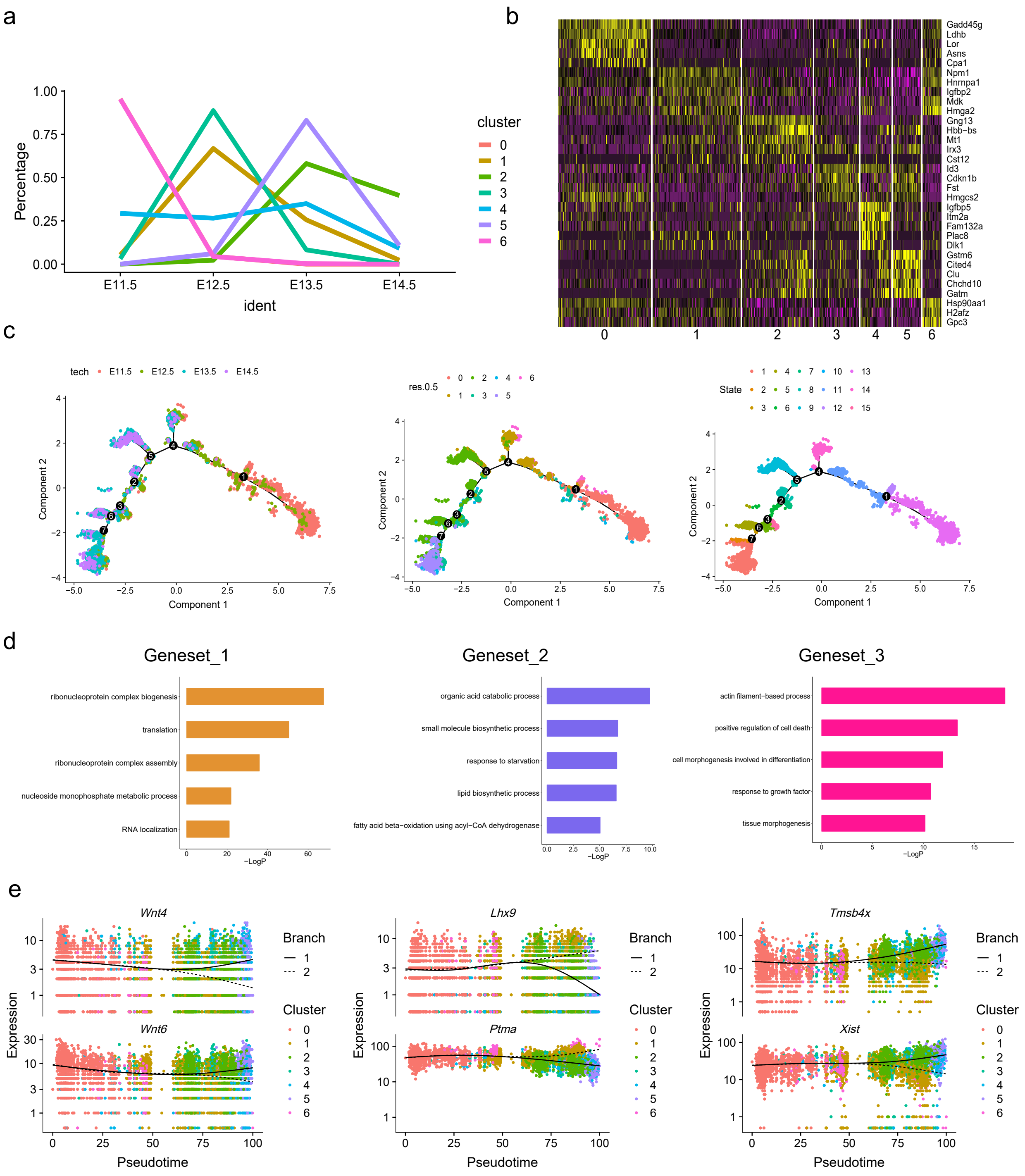

### Supplementary Figure 7

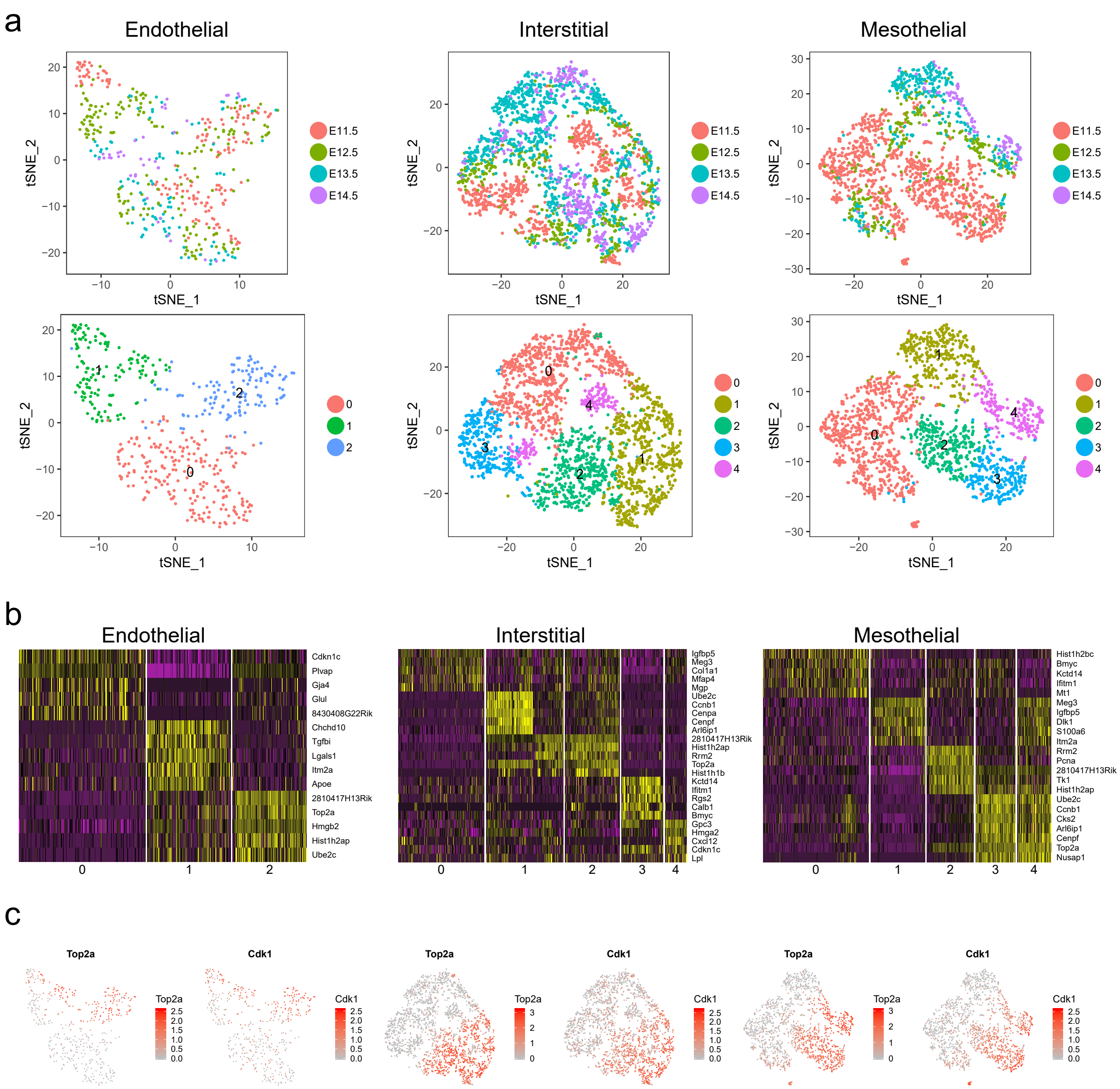
